# Mapping lesion, structural disconnection, and functional disconnection to symptoms in semantic aphasia

**DOI:** 10.1101/2021.12.01.470605

**Authors:** Nicholas E. Souter, Xiuyi Wang, Hannah Thompson, Katya Krieger-Redwood, Ajay D. Halai, Matthew A. Lambon Ralph, Michel Thiebaut de Schotten, Elizabeth Jefferies

## Abstract

Patients with semantic aphasia have impaired control of semantic retrieval, often accompanied by executive dysfunction following left hemisphere stroke. Many but not all of these patients have damage to the left inferior frontal gyrus, important for semantic and cognitive control. Yet semantic and cognitive control networks are highly distributed, including posterior as well as anterior components. Accordingly, semantic aphasia might not only reflect local damage but also white matter structural and functional disconnection. Here we characterise the lesions and predicted patterns of structural and functional disconnection in individuals with semantic aphasia and relate these effects to semantic and executive impairment. Impaired semantic cognition was associated with infarction in distributed left-hemisphere regions, including in the left anterior inferior frontal and posterior temporal cortex. Lesions were associated with executive dysfunction within a set of adjacent but distinct left frontoparietal clusters. Performance on executive tasks was also associated with interhemispheric structural disconnection across the corpus callosum. In contrast, poor semantic cognition was associated with small left-lateralized structurally disconnected clusters, including in the left posterior temporal cortex. Little insight was gained from functional disconnection symptom mapping. These results demonstrate that while left-lateralized semantic and executive control regions are often damaged together in stroke aphasia, these deficits are associated with distinct patterns of structural disconnection, consistent with the bilateral nature of executive control and the left-lateralized yet distributed semantic control network.

## 1. Introduction

To understand the world around us, we draw on two connected but dissociable components; a store of long-term semantic knowledge (e.g., Patterson et al., 2007) and control processes that shape retrieval to suit current circumstances (Lambon Ralph et al., 2017; Rogers et al., 2015). This distinction is highlighted by comparing semantic dementia (SD) and semantic aphasia (SA).

SD patients show conceptual degradation associated with atrophy of the ventral anterior temporal lobes (vATL) and highly consistent semantic deficits across tasks that probe the same items (Bozeat et al., 2000; Mummery et al., 2000), in line with the view this site acts as a ‘semantic hub’ for heteromodal concepts (Lambon Ralph et al., 2017; Patterson et al., 2007). According to the hub-and-spoke model of semantic cognition, the vATL ‘hub’ works in concert with modality-specific ‘spokes’ in order to generate generalisable and coherent representations (Rogers et al., 2021). In contrast to those with SD, patients with SA have intact conceptual representations but an impaired ability to retrieve information in a flexible and context-appropriate manner, following left inferior frontal and/or temporoparietal stroke (Jefferies, 2013; Jefferies & Lambon Ralph, 2006)^1^. Such deficits are multi-modal, such that these patients experience impairments in non-verbal attribution of appropriate object use (Corbett et al., 2011), as well as in verbal semantic matching between words (Noonan et al., 2010). SA patients show stronger-than-normal effects of cues and miscues designed to help or hinder conceptual retrieval (Jefferies et al., 2008; Lanzoni et al., 2019; Noonan et al., 2010). These effects are accompanied by poor retrieval of weak associations and the subordinate meanings of ambiguous words and objects, as well as more significant impairment when targets are presented alongside strong distractors (Noonan et al., 2010).

Consequently, the study of these patients can provide insights into the neurocognitive mechanisms that allow us to use our conceptual knowledge in a controlled way.

Semantic deficits in SA are thought to reflect damage to a distributed but largely left-lateralised semantic control network (SCN), but evidence demonstrating the functional relevance of connectivity between key SCN nodes is lacking. While the peak lesion overlap in SA is in the left posterior inferior frontal gyrus (IFG), not every case shows damage here.

Lesions are also highly variable within the left parietal and posterior temporal cortex (Chapman et al., 2020; Hallam et al., 2018; Lanzoni et al., 2019; Stampacchia et al., 2018). This lesion heterogeneity is anticipated by neuroimaging meta-analyses of healthy participants (Jackson, 2021; Noonan et al., 2013), which highlight reliable activation of posterior components of the SCN, most notably the left posterior middle temporal gyrus (pMTG), along with the IFG in tasks with high semantic control demands (e.g., Becker et al., 2020; Krieger-Redwood et al., 2015; Zhang et al., 2021). Studies employing inhibitory stimulation suggest that both the left IFG and pMTG play a critical role in semantic control (Davey et al., 2015; Whitney et al., 2011). Moreover, damage or inhibitory stimulation of the left IFG elicits increased activation in the left pMTG (Hallam et al., 2018; 2016), as would be expected within a distributed functional network. While this work suggests that both anterior and posterior sites support semantic control, the structural and functional disconnection that is anticipated from the diverse lesions in SA has not been clearly delineated. We might expect similar and overlapping disconnection patterns across cases with different lesions (affecting anterior and posterior components of the SCN, for example), given the same network is thought to be damaged in patients with semantic deficits.

Another unresolved issue concerns the extent to which the SCN is distinct from the multiple-demand network (MDN), which supports executive control across domains (Fedorenko et al., 2013). Like the SCN, the MDN includes highly distributed anterior and posterior components and neuroimaging studies of healthy participants suggest these networks are adjacent in lateral temporal and lateral and medial frontal cortex (Davey et al., 2016; Gao et al., 2021; Wang et al., 2020). This proximity of the SCN and MDN in the left hemisphere may explain why SA patients, selected to show heteromodal semantic deficits, frequently also present with non-semantic executive impairment (Thompson et al., 2018).

Since MDN regions support the performance of semantic tasks with high control demands (Krieger-Redwood et al., 2015; Wang et al., 2020), damage to these regions will contribute to difficulties regulating semantic cognition. Nevertheless, the SCN and MDN diverge in their degrees of lateralisation. While the SCN is largely left-lateralised, the MDN comprises distributed bilateral regions (Camilleri et al., 2018; Gao et al., 2021; Gonzalez Alam et al., 2019). As such, effective semantic control should involve connectivity within the left-hemisphere, while domain-general control should rely more on interhemispheric connectivity (Gonzalez Alam et al., 2021), allowing the integration of information across right-hemisphere regions dominant in the control of visuospatial processing with contralateral frontal regions (Wu et al., 2016). As a result, structural or functional disconnection-symptom mapping may separate semantic control impairment from general executive deficits in a way that cannot be achieved by lesion-symptom mapping in patients with left-hemisphere stroke.

Unlike the MDN, the SCN also overlaps with regions of the default-mode network (DMN) – indeed, nodes of the SCN fall in between MDN and DMN in the left hemisphere (Davey et al., 2016; Gao et al., 2021; Wang et al., 2020). Posterior aspects of the left IFG, bordering the inferior frontal sulcus, overlap with MDN and are implicated in executive control, while anterior aspects of the left IFG that fall within the DMN are thought to be engaged more specifically in controlled semantic retrieval (Badre & Wagner, 2007; Krieger-Redwood et al., 2015; Zhang et al., 2021). The DMN also extends beyond semantically relevant regions; this large bilateral network is associated with task-related deactivation (Raichle, 2015), the coordination of information across the cortex (Lanzoni et al., 2020) and multiple forms of abstract, internal and heteromodal cognition (Gordon et al., 2020; Murphy et al., 2018; Smallwood et al., 2021). In addition to overlapping with SCN in left lateral posterior temporal and ventral and dorsomedial prefrontal cortex (Davey et al., 2016; Gao et al., 2021; Wang et al., 2020), DMN overlaps with key heteromodal sites relevant for semantic cognition irrespective of control demands, including semantic regions in the ATL (Smallwood et al., 2021) and angular gyrus (Vatansever et al., 2017).

Given the above evidence, we might expect semantic and executive deficits in SA following left-hemisphere stroke to be associated with similar lesion profiles but distinct patterns of disconnection. Structural and functional disconnection within the left-lateralised components of the SCN may underpin semantic deficits. At the same time, broader executive impairment across domains may relate to disconnection between left and right control regions. Studies already show that white matter connections between anterior temporal and occipital/middle temporal regions predict semantic impairment beyond the contribution of grey matter damage alone (Fang et al., 2018). Tracts implicated in semantic cognition in the left hemisphere include the inferior fronto-occipital fasciculus (IFOF), anterior thalamic radiation (ATR), uncinate fasciculus (UF) and inferior longitudinal fasciculus (ILF; Almairac et al., 2015; Han et al., 2013; Sierpowska et al., 2019). The left ILF and left IFOF have been associated with semantic control specifically (Marino et al., 2020; Nugiel et al., 2016).

Moreover, distinct changes in structural connectivity are associated with semantic impairment in SD and SA patients: SA is related to changes in left frontal-subcortical and left frontal-temporal/occipital networks, while symptoms in SD are associated with fractional anisotropy of a left medial temporal white matter network (Ding et al., 2020). Conversely, a decline in executive function occurs with compromised integrity of the corpus callosum, in both healthy ageing (Johnson et al., 2017; Jokinen et al., 2007; Voineskos et al., 2012) and patient groups (Bodini et al., 2013), consistent with the theory that demanding tasks rely on cross-hemispheric integration (Gazzaniga, 2005; Schulte & Müller-Oehring, 2010).

Accordingly, impairments in semantic control and executive function may be predicted by left-lateralised and bilateral disconnection, respectively.

This study aimed to characterise typical patterns of infarct, plus structural and functional disconnection, in a sample of 23 SA patients. The use of such ‘connectomic’ data may help to elucidate the relationship between diffuse networks and impairments following cerebral insult (Fornito et al., 2015). We assessed the impact of lesion on functional networks including the SCN, MDN and DMN, given all these networks are implicated in semantic cognition. Individual patterns of structural disconnection were predicted by tracking white matter fibres likely to pass through a patient’s lesion based on diffusion-weighted imaging data from neurologically healthy participants (Foulon et al., 2018). Structural disconnection in stroke patients assessed in this way has been shown to reflect functional activation better than lesion location alone (Thiebaut de Schotten et al., 2020), and to predict post-stroke symptoms including apraxia (Garcea et al., 2020) and executive impairment (Langen et al., 2018). Similarly, measures of functional disconnection were derived by assessing patterns of intrinsic functional connectivity with the lesion site, using resting-state scans from neurologically healthy participants and seed-based functional connectivity analysis (Salvalaggio et al., 2020). Although this functional disconnection metric has been shown to be less predictive of cognitive deficits than structural disconnection (Salvalaggio et al., 2020), here we implemented a method thought to have higher sensitivity, which involved performing principal components analysis of resting-state data from a sample of healthy adults from each patient’s lesion and seeding the first component of this functional connectivity pattern. Evidence from Pini et al. (2021) suggests that this approach provides better anatomical specificity and behavioural prediction than seeding the entire lesion.

We explored associations between measures of lesion location, structural disconnection, and functional disconnection and performance on semantic cognition and non-semantic executive function tests. We would expect adjacent or similar lesions to predict deficits of semantic cognition and executive control, given that the networks supporting these functions in the left hemisphere are thought to have a similar topological organisation.

Moreover, we would expect *peak* lesion location to fall within the SCN, given that SA patients show deregulated conceptual retrieval. We may also predict some lesion extension into adjacent DMN, MDN, and core semantic regions, given their spatial proximity to the SCN and potential contributions to semantic cognition. These lesions should be accompanied by widespread structural and functional disconnection that is again maximal within the SCN. However, since executive control is thought to draw on the bilateral MDN, while semantic control is strongly left-lateralised, the broader patterns of disconnection that predict performance within these domains may be more distinct. Accordingly, poor semantic cognition should be associated with left-lateralised patterns of structural and functional disconnection (across left-hemisphere components of DMN, SCN and MDN). Executive dysfunction on non-semantic tasks might be related to more bilateral patterns of disconnection specifically within MDN.

## 2. Method

### 2.1. Participants

Participants were 23 SA patients with impaired multimodal semantic control following left hemisphere stroke. Patients were recruited from communication support groups across Yorkshire, Surrey, and Manchester in the UK. Patients were all right-handed and native English speakers, had a mean age of 62.2 (SD = 11.9) and a mean age of leaving education of 16.3 (SD = 1.5)^2^. Participants underwent structural MRI and cognitive testing at least six months post-stroke, with a mean of 6.7 years (SD = 5.5) since the infarct, such that patients were in the chronic phase and changes in brain function would have been relatively gradual and subtle. Scans were obtained close in time to the behavioural testing. Volunteers were excluded if they reported participating in ongoing rehabilitation at the time of testing.

All patients reported no history of traumatic brain injury or suspected or confirmed neurodegenerative disorders. Patients were selected to show impairment on at least one verbal and one nonverbal measure of semantic control, in line with Head’s (1926) definition of SA as multimodal impairment in the manipulation of knowledge for symbolic processing.

### 2.2. Background neuropsychological testing

Patients completed a series of tests probing language, memory, executive function, visuospatial processing, and semantic cognition. Patients’ individual performance on tests of background neuropsychology and semantic cognition can be seen in Supplementary Table 1 and Supplementary Table 2, respectively. Patients had variable impairment of word repetition and commonly showed impaired category fluency, letter fluency and verbal working memory. Fourteen patients were impaired on at least one test of executive function. Most patients had relatively good visuospatial processing. A description of these tasks and performance is provided in Supplementary Materials (*Background Neuropsychology*).

All patients were impaired on at least one verbal and one nonverbal measure of semantic cognition, in line with our inclusion criteria. We found 78% of patients were impaired on the word version of Camel and Cactus Test (CCT) of semantic associations, while 52% were impaired on the picture version. Most patients showed near-ceiling performance on a simple test of word-picture matching, as has previously been shown (e.g., Lanzoni et al., 2019; Thompson et al., 2018). The majority of cases (80%) showed impaired picture naming. All patients who were tested with phonemic cueing showed evidence of improved accuracy as a result, except for patients performing at floor level. Similarly, when the relevant data was available, patients showed strong effects of semantic control manipulations across a series of assessments, including difficulty retrieving subordinate conceptual information, susceptibility to contextual cues and miscues, and deleterious effects of semantic distractors on synonym judgement.

Principal components analysis (PCA) with oblique rotation was used to extract a composite score for semantic tasks that were maximally available across the sample: namely word and picture CCT, and overall performance on the no cue version of the ‘ambiguity’ assessment (Noonan et al., 2010). In doing so, we aimed to obtain a single score to reflect semantic performance across tasks for each participant. PCA was performed with data from the 21 patients who completed all three tests. This analysis revealed that all three tests loaded strongly on a single component with an eigenvalue of 2.6 (loadings: CCT words = .95; CCT pictures = .92; ambiguity = .92) that explained 87% of variance in performance. We extracted a ‘semantic cognition composite score’ from this component, with lower values reflecting greater impairment of semantic cognition. Patients’ individual composite scores are presented in Supplementary Table 2. Performance on the Brixton Spatial Anticipation Test (Burgess & Shallice, 1997) was taken to reflect patients’ degree of impairment in cognitive control beyond the semantic domain^3^, for the 20 patients for whom data were available. This task involved anticipating the locations where a dot would move to, given past patterns, and shifting these predictions when the pattern changed. Performance on the Brixton test correlated positively with patients’ semantic composite scores (*r*(18) = .61, *p* = .005), providing evidence that poorer semantic performance was accompanied by poorer executive control on a non-verbal task.

In summary, while SA patients were screened in on the basis of verbal and nonverbal semantic impairment, they were not excluded on the basis of impairment in other domains. Many SA patients in this sample presented with multi-domain impairment beyond semantic cognition, in common with earlier studies (e.g., Thompson et al., 2018). The symptom mapping employed in the current investigation can therefore test for dissociable substrates underlying semantic and domain-general cognitive control impairments.

### 2.3. MRI acquisition

Structural T1 images were obtained for all patients. Patients in York (N = 13) were scanned using a 3T GE HDx Excite MRI scanner on a T1-weighted 3D fast spoiled gradient echo sequence (TR = 7.8ms, TE = minimum full, flip-angle = 20°, matrix size = 256 x 256, 176 slices, voxel size = 1.13 x 1.13 x 1mm). Patients in Manchester (N = 8) were scanned using a 3T Philips Achieva MRI scanner using a T1-weighted 3D inversion recovery sequence (TR = 9.0ms, TE = 3.93ms, flip-angle = 8°, matrix size = 256 x 256, 150 slices, voxel size = 1 x 1 x 1mm). Scanning parameters for patients scanned in Surrey (N = 2) are not available.

For analysis of functional disconnection, we used resting state scans from an independent sample of 207 neurologically healthy volunteers recruited from the University of York. Data were collected from the whole brain on a 3T GE HDx Excite MRI scanner using single-shot 2D gradient-echo-planar imaging (TR = 3s, TE = minimum full, flip-angle = 90°, matrix size = 64 x 64, 60 slices, voxel size = 3 x 3 x 3mm, 180 volumes). We excluded sixteen participants: nine due to missing behavioural data, one due to missing MRI data, one due to incorrect TR in MRI acquisition, and four during pre-processing because they exceeded our motion cut-off of .3 mm, had more than 20% invalid scans and/or mean global signal change of z > 2. The final sample size, therefore, consisted of 191 participants.

Estimates of structural disconnection were based on diffusion-weighted data from healthy controls, collected on a 3T GE Signa HDx TwinSpeed system (TR = 20/30 R-R intervals, TE = 93.4ms, matrix size = 128 x 128, 60 slices, voxel size = 2.4 x 2.4 x 2.4mm).

### 2.4. Lesion segmentation

Patients’ T1 scans underwent brain extraction in ANTs using a template from the OASIS Brain Project (https://www.oasis-brains.org/; Marcus et al., 2010). Registration to MNI152 space was also performed using ANTs linear registration (version 2.1.0; Avants et al., 2011; 2014), which utilises a symmetric normalisation model. Default parameters were used including aligning centres and orientations, accounting for scaling factors, ending with affine transformation, and including optimisation for cost function (Avants et al., 2011).

Patients’ lesions were then manually drawn in MRICron, using a combination of the 3D fill tool feature and subsequent validation of each slice. Care was taken to avoid implicating enlarged sulci or ventricles in the lesion: cases where sulci or ventricles have been implicated by the 3D tool are typically observable through visual inspection by cross-referencing axial, sagittal, and coronal views of a given slice, and in such cases the respective highlighting was removed from the lesion drawing.

### 2.5. Structural disconnection

The BCBtoolkit (Foulon et al. 2018; http://www.toolkit.bcblab.com) was used to generate probabilistic maps of spreading structural disconnection. This approach uses a set of ten healthy controls’ DWI datasets (Rojkova et al., 2016) to track fibers passing through each patient’s lesion. This makes it possible to estimate likely structural disconnection from a lesion even when no diffusion-weighted imaging is acquired. The control sample used here is independent of the sample used to estimate functional disconnection (see Section 2.3).

Patients’ lesions in the MNI152 are used as seeds for tractography in Trackvis (Wang et al., 2007). Tractographies from the lesions were transformed in visitation maps (Thiebaut de Schotten et al., 2011) and binarised. Finally, a percentage overlap map was produced by summing the normalized visitation map of each healthy subject at each point in MNI space. In the resulting disconnectome map for each patient, the value of each voxel reflects the interindividual variability of tract reconstructions in controls, resulting in a value between 0 and 1 reflecting the probability of disconnection (Thiebaut de Schotten et al., 2015).

Disconnectome maps for each patient were thresholded at 0.5, such that the disconnectome maps corresponded to the exact white matter anatomy of more than 50% of the healthy controls (Foulon et al., 2018). While 50% is the default threshold in the BCBtoolkit, disconnectome maps were also generated at thresholds of 40% and 60% to test the stability of our analysis. Symptom mapping at these alternative thresholds (using the same procedure outlined in section 2.8) produced similar clusters to those identified at the 50% threshold, as seen in Supplementary Figure 1.

In order to assess the effect of structural disconnection on specific white matter tracts, probabilistic tracts included in the BCBtoolkit (Foulon et al., 2018) were extracted, thresholded at a value of 0.95, and used to visualise the tracts of interest. We were specifically interested in tracts implicated in semantic cognition, language, domain-general cognitive control, or the confluence of these functions (Agosta et al., 2010; Almairac et al., 2015; Bodini et al., 213; Dick et al., 2019; Ding et al., 2020; Han et al., 2013; Huang et al., 2015; Johnson et al., 2017; Marino et al., 2020; Nugiel et al., 2016; Rizio & Diaz, 2016; Sierpowska et al., 2019; Spitz et al., 2013), including the uncinate fasciculus (UF), anterior thalamic radiation (ATR), inferior longitudinal fasciculus (ILF), inferior fronto-occipital fasciculus (IFOF), frontal aslant tract (FAT), arcuate fasciculus (AF), superior longitudinal fasciculus (SLF), and corpus callosum. We quantitatively assessed the effect of structural disconnection on these tracts using the Tractotron function of the BCBtoolkit (Foulon et al., 2018). Tractotron maps the lesion from each patient onto tractography reconstructions of specific white matter pathways obtained from a group of healthy controls (Rojkova et al., 2016), and quantifies both the probability of a given tract being disconnected, and the proportion of this tract that is likely to be disconnected (Thiebaut de Schotten et al. 2014).

### 2.6. Functional disconnection

We generated maps of functional connectivity to each patient’s lesion location, to provide an indirect measure of functional disconnection. Typically, functional disconnection has been estimated by using patients’ entire binarised lesion in seed-based functional connectivity analysis (e.g., Salvalaggio et al., 2020). However, this approach can be problematic when seeding large lesions, which contain multiple functionally distinct regions that couple to different networks (Boes, 2021; Bowren et al., 2022). We therefore adopted a method proposed by Pini et al. (2021), whereby PCA is first conducted on the connectivity of all voxels within each lesion in a sample of neurologically healthy adults (here, the same 191 participants described in Section 2.3). Forty components were extracted for each lesion.

Given that the first principal component extracted explains the largest amount of variance, this component was taken as the main within-lesion connectivity axis. The first component for each patient was thresholded such that only voxels within the lesion with an absolute coefficient above the 20^th^ percentile were retained. This threshold has been shown to provide higher functional specificity than higher and lower percentile thresholds (Pini et al., 2021).

The resulting thresholded map was then binarised and seeded in functional connectivity analysis using an independent sample of resting-state fMRI scans from neurologically healthy participants (as described in Section 2.3). Compared with seeding the entire lesion, this approach has been shown to improve behavioural prediction and anatomical specificity of estimates of functional disconnection (Pini et al., 2021). All PCAs and functional connectivity analyses were conducted using the CONN functional connectivity toolbox V.20.b (Whitfield-Gabrieli & Nieto-Castanon, 2012; www.nitrc.org/projects/conn) in MATLAB.

Functional resting-state volumes were skull-stripped, slice-time (interleaved) and motion-corrected, and co-registered to the high-resolution structural image, spatially normalised to MNI space using the unified-segmentation algorithm, smoothed with a 6 mm FWHM Gaussian kernel, and band-passed filtered (.008-.09 Hz) to reduce low-frequency drift and noise effects. A pre-processing pipeline of nuisance regression included motion (twelve parameters: the six translation and rotation parameters and their temporal derivatives) and scrubbing (all outlier volumes were identified through the artifact detection algorithm included in CONN with conservative settings: scans for each participant were flagged as outliers based on scan-by-scan change in global signal above z > 3, subject motion threshold above 5 mm, differential motion and composite motion exceeding 95% percentile in the normative sample). We used the anatomical CompCor approach, a PCA-based approach which attempts to isolate aspects of images caused by artefacts (Satterthwaite et al., 2019), and removes these nuisance variables in a single linear regression step in order to provide a clear signal (Behzadi et al., 2007). CompCor also includes a linear detrending term, eliminating the need for global signal normalisation. A covariate containing quality assurance parameters flagged average subject motion as outliers for scrubbing at a threshold of .5 mm based on framewise displacement. Group-level analyses in CONN were cluster-size FWE corrected and controlled for the number of seeds (Bonferroni, p < .002), and used a height threshold of p < .001. The resulting output files were thresholded such that only positively associated voxels remained and were taken to reflect patients’ maps of functional disconnection.

### 2.7. Functional networks

We assessed the effect of patients’ lesions on networks of interest. To identify the SCN, we used the map from Jackson (2021), derived from a meta-analysis of studies that manipulated semantic control demands. To identify semantic regions outside the SCN, we used a map from the meta-analytic tool Neurosynth (Yarkoni et al., 2011) for the term ‘Semantic’. The MDN was defined using a map from Fedorenko et al. (2013) reflecting activation associated with global effects of difficulty across seven diverse tasks, thresholded at t > 1.5. We also identified areas common to both the SCN and MDN, using a conjunction of the respective maps described above. Finally, the DMN was defined using the 7-network parcellation from Yeo et al. (2011). All maps were mutually exclusive such that (1) any voxels contained within the MDN, SCN, or Neurosynth semantic network were removed from the DMN (i.e. this map focussed on domain-general responses, excluding regions specifically implicated in semantic cognition); (2) any voxels contained within the SCN or MDN were removed from the Neurosynth semantic network (i.e., this map excluded both DMN and control regions and as such focussed on those involved in semantic representation or more automatic aspects of retrieval and not more controlled patterns of retrieval); and (3) any voxels contained within both the SCN and MDN were removed from each individual map and placed in the conjunctive ‘MDN + SCN’ map (such that the SCN-only regions were not implicated in domain-general control). Visualisations of each mutually exclusive network can be seen in Figure 2 when results of the network damage analysis are reported.

We assessed the mean percentage of each network that was lesioned across the sample. As all lesions were restricted to the left hemisphere, each network was confined to the left hemisphere for this analysis. All network maps were binarized prior to analysis. We identified for a given patient, the number of voxels implicated in both their binary lesion file and a given functional network, computed as a percentage of the total number of voxels implicated in this network. This process was also conducted for patients’ functional disconnection maps (Supplementary Figure 2). We did not assess overlap with the structural disconnection maps since these were confined to white matter.

The SCN map (Jackson, 2021) spans five distinct clusters including (1) left frontal regions (left IFG, insula, orbitofrontal cortex, and precentral gyrus), (2) left posterior temporal regions (left pMTG, posterior inferior temporal gyrus, and posterior fusiform gyrus), (3) the bilateral dmPFC, (4) the right IFG (pars orbitalis) and insula, and (5) the right IFG (pars triangularis). The SCN was split into these separate clusters in order to observe spreading disconnection within the SCN beyond lesion site. We extracted the mean percentage of each cluster that overlapped with each patient’s lesion map; we then identified whether patients’ structural and functional disconnection maps showed any overlap with each SCN cluster that fell outside the patients’ lesion.

### 2.8. Symptom mapping

We assessed the patterns of lesion, structural disconnection, and functional disconnection associated with the semantic composite score and executive function performance. Patients’ binary lesion segmentations, continuous structural disconnection maps, and continuous functional disconnection maps were separately entered into nonparametric 2-sample t-tests in Randomise (Winkler et al., 2014; https://fsl.fmrib.ox.ac.uk/fsl/fslwiki/Randomise/UserGuide), using 5,000 permutations. Threshold-free cluster enhancement was implemented to identify clusters (Smith & Nichols, 2009). This analysis was restricted to the 20 patients for whom both the semantic cognition composite and Brixton test scores were available. Each model was set up such that the input was a 4D file containing all patient maps. In each case simultaneous regressors included the size of a given patient’s respective input file size (i.e., the overall size of the binary lesion map or disconnection map), their semantic composite, and their Brixton performance. Within each model, clusters implicated for a given behavioural measure therefore regressed out the size of the input file as well as performance on the other behavioural measure. A binary mask containing the addition of all patient input files was used. Contrasts were specified such that the resulting output revealed (i) clusters associated with better semantic performance, (ii) clusters associated with poorer semantic performance, (iii) clusters associated with better Brixton performance, and (iv) clusters associated with poorer Brixton performance. All resulting t-value maps were thresholded at a value of 2.6 for interpretation.

Lesioned/disconnected clusters associated with better behavioural performance reflect better performance relative to the sample mean, rather than absolute improvements in performance as a result of damage. Such clusters were not of explicit interest in the current analysis, so are reported in Supplementary Figure 3.

## 3. Results

Data for this project are available on the Open Science Framework (https://osf.io/6psqj/). Unthresholded group-level NIFTI files corresponding to the results presented here can be seen on Neurovault (https://neurovault.org/collections/10333/).

### 3.1. Lesion profile

We first characterised the regions most typically lesioned across the sample. The lesion group map reflecting maximum overlap is provided in Figure 1a. An unthresholded view provided in Supplementary Figure 4a. Due to the relative heterogeneity of the lesions (see Figure 1d), this image has a minimum threshold of four cases, while a higher minimum threshold of 19 cases is used for the structural and functional disconnection overlap maps. Lesions encompassed a range of left frontoparietal regions; for each anatomical region extracted from the Harvard-Oxford atlas^4^, we calculated the percentage of the sample (N = 23) that showed some evidence of damage. These regions included the left IFG (pars triangularis, 56.5%, and pars opercularis, 69.6%), frontal pole (52.2%, with this region including pars orbitalis), middle frontal gyrus (MFG; 60.9%), insular cortex (60.9%), precentral gyrus (82.6%), postcentral gyrus (73.9%), supramarginal gyrus (SG; 65.2%), and angular gyrus (AG; 65.2%). In some cases, lesions extended to temporal and occipital regions, including the pMTG (34.8%), temporo-occipital part of the MTG (39.1%), superior temporal gyrus (STG; 47.8%), inferior temporal gyrus (ITG; 34.8%), planum temporale (52.2%), temporal pole (43.5%), lateral occipital cortex (LOC; 69.6%), and occipital pole (30.4%). Damage to the temporal pole spared vATL in every case. Some of the damaged described here is not visible in Figure 1a due to the heterogeneity of lesion location. For example, while eight patients show damage to some aspect of the pMTG, this damage does not always fall in the same voxels, meaning this region does not appear to be impacted to this extent in the thresholded overlap.

**Figure 1.**
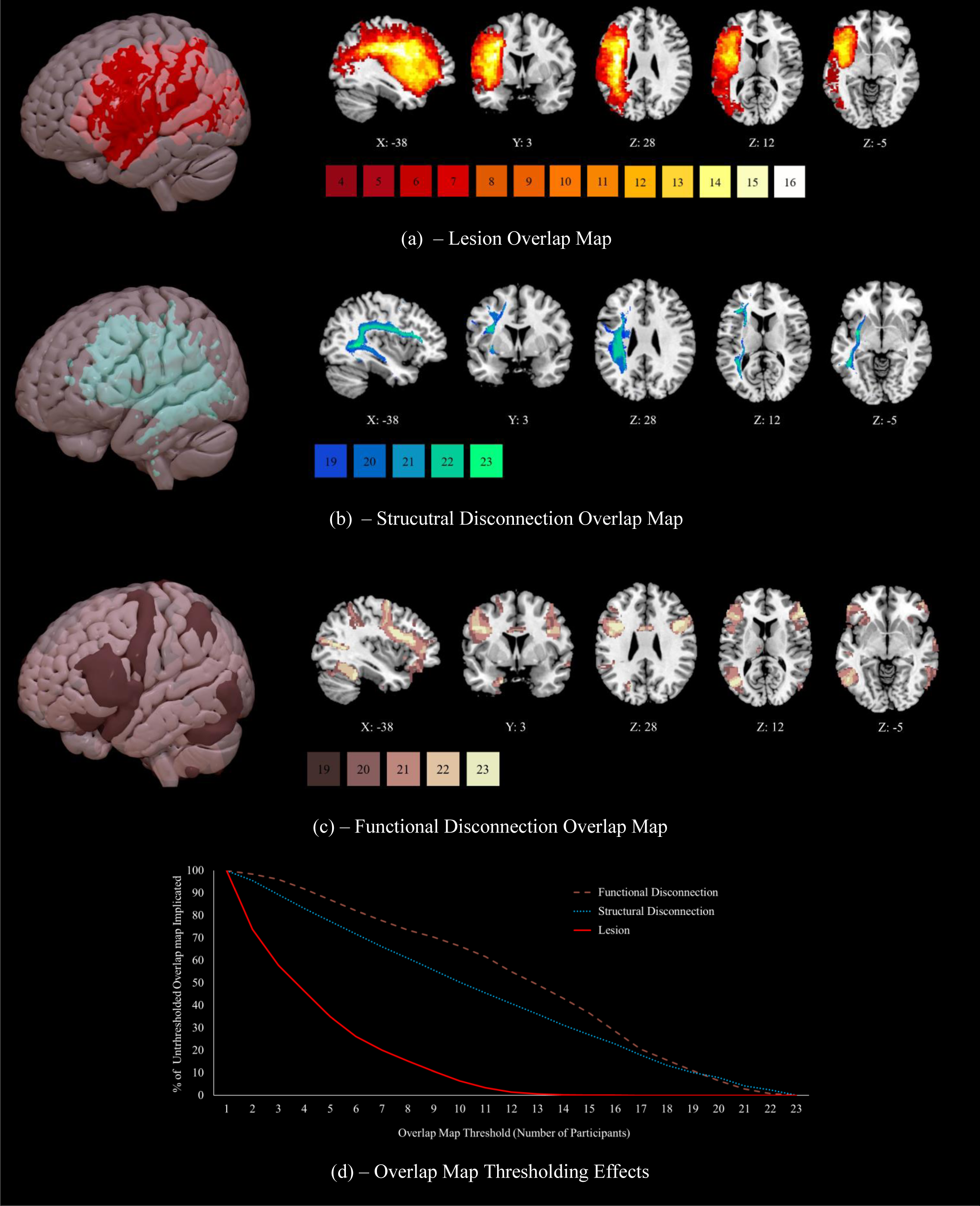
Maximum overlap for (a) lesion sites thresholded at 4 cases, (b) structural disconnection maps, generated using the BCB Toolkit, thresholded at 19 cases, and (c) functional disconnection maps, generated using CONN, thresholded at 19 cases. 3D rendering generated in SurfIce. (d) Visualisation of the numbers of cases showing overlapping lesions, structural disconnection (confined to white matter) and functional disconnection (confined to grey matter), expressed as a percentage of the total number of voxels damaged or disconnected for at least one participant. This figure demonstrates the homogeneity of patterns of anticipated structural and functional disconnection, relative to the lesion overlap map. N = 23. DESCRIPTIVE CAPTION FOR TEXT-TO-SPEECH: Figure 1 presents overlap maps for the measures of lesion, structural disconnection, and functional disconnection. Figure 1a presents lesion overlap between 4 and 16 cases. Lesions are confined to the left hemisphere and are frontoparietal, peaking in the precentral and middle frontal gyri. Figure 1b presents structural disconnection overlap between 19 and 23 cases. Structural disconnection is left lateralised at this threshold and shows maximal overlap with the superior longitudinal fasciculus and inferior fronto-occipital fasciculus. Figure 1c presents functional disconnection overlap between 19 and 23 cases. Functional disconnection is bilateral and extensive, peaking in the temporooccipital part of the left inferior temporal gyrus and the right inferior frontal gyrus pars triangularis. A line graph in Figure 1d reflects the homogeneity of structural and functional disconnection relative to lesion location, with increasing thresholds reducing the overall size of the lesion overlap map at a much higher rate than for either disconnection measure.

### 3.2. Structural disconnection

Using patients’ lesions in conjunction with the disconnectome function of the BCB Toolkit, we extracted maps of probabilistic spreading white matter structural disconnection for each patient in order to characterise typical patterns of diffuse damage beyond lesion site. The structural disconnection group map reflecting maximum overlap can be seen in Figure 1b. An unthresholded view provided in Supplementary Figure 4b. Extensive structural disconnection was found to be likely throughout the left hemisphere. Twenty-one patients showed some evidence of right hemisphere structural disconnection. While disconnection in the left hemisphere was convergent, right hemisphere disconnection was relatively heterogenous, meaning it cannot be observed in Figure 1b at the selected threshold of 19 cases (observable at threshold of 17, see Figure S4b). Supplementary Figure 5 shows the mean probability of each tract being disconnected (Figure S5a), as well as estimates of the mean proportion of each tract that was disconnected across the sample (Figure S5b), both estimated using Tractotron (see Section 2.5). This figure also provides visualisations of the structural disconnection overlap map (Figure S5c) and each tract of interest (Figure S5d-k). The mean probability of disconnection was above 60% for all tracts that were examined. The mean proportion of disconnection was highest for ILF, SLF III and anterior AF. Figure 1d suggests that the patterns of structural disconnection seen in this SA patient sample were relatively homogeneous and highly overlapping, despite their heterogenous lesions. Applying a threshold of ten patients reduces the size (in voxels) of the overall lesion overlap map by 93.5%, while this same threshold reduces the structural disconnection overlap map by only 49.7%.

### 3.3. Functional disconnection

In order to assess the intrinsic functional architecture associated with structural disconnection, we subjected each patient’s lesion to PCA, and treated the first component as the main within-lesion connectivity axis, which was then used a seed in a resting-state analysis of an independent sample of healthy individuals in CONN to provide maps of functional disconnection. Figure 1c provides the functional disconnection group map reflecting maximum overlap. An unthresholded view is provided in Supplementary Figure 4c. The resting-state maps associated with all 23 patients’ lesions showed functional disconnection bilaterally of the frontal pole (including pars orbitalis), IFG (pars opercularis and pars triangularis), insular cortex, MFG, precentral gyrus, postcentral gyrus, and posterior SG, and of the right AG, left posterior ITG, left inferior LOC, right superior LOC, and left temporooccipital parts of the MTG and ITG. In most cases, patients’ functional disconnection maps predicted disconnection of the superior frontal gyrus (SFG; 95.7%)^5^, pMTG (95.7%), aMTG (left = 73.9%, right = 65.2%), anterior ITG (left = 73.9%, right = 82.6%), anterior STG (left = 87.0%, right = 91.3%), posterior STG (left = 91.3%, right = 95.7%), anterior SG (left = 95.7%, right = 91.3%), occipital pole (91.3%), and planum temporale (73.9%), and of the left AG (95.7%), right inferior LOC (95.7%), left superior LOC (95.7%), right posterior ITG (95.7%), and right temporooccipital parts of the ITG and MTG (95.7%). These patterns of functional disconnection were relatively homogeneous, with a threshold of ten patients reducing the overall size of the functional disconnection overlap map by only 33.6% (see Figure 1d).

### 3.4. Damage and disconnection within functional networks

#### 3.4.1. Mean percent lesioned

We observed the extent of lesion to each functional network of interest. The mean percentage of voxels lesioned in left hemisphere aspects of DMN, semantic non-control, SCN, SCN+MDN and MDN networks is shown in Figure 2a^6^. The greatest damage was seen in SCN regions (Figure 2d), followed by SCN+MDN (Figure 2e). The MDN (Figure 2f) and non-control semantic regions (Figure 2c) showed more modest damage, and damage to the DMN was minimal (Figure 2b). An equivalent analysis of functional disconnection is provided in Supplementary Figure 2. Most (≥ 57.8%) of the SCN, MDN, SCN+MDN and core semantic regions were functionally disconnected across the sample. The DMN was also affected but across less than half of the network (30.8%). As seen in Supplementary Table 3, a significantly smaller percentage of the DMN was lesioned than other functional networks, excluding core semantic regions. The extent of lesion did not differ significantly between other networks of interest. A significantly smaller percentage of the DMN was also functionally disconnected when compared to all other networks. Furthermore, a greater percentage of voxels in the SCN and shared between the SCN and MDN were functionally disconnected than in core semantic regions. A greater percentage of voxels shared between the SCN and MDN were functionally disconnected when compared to the MDN alone (see Supplementary Table 3).

**Figure 2.**
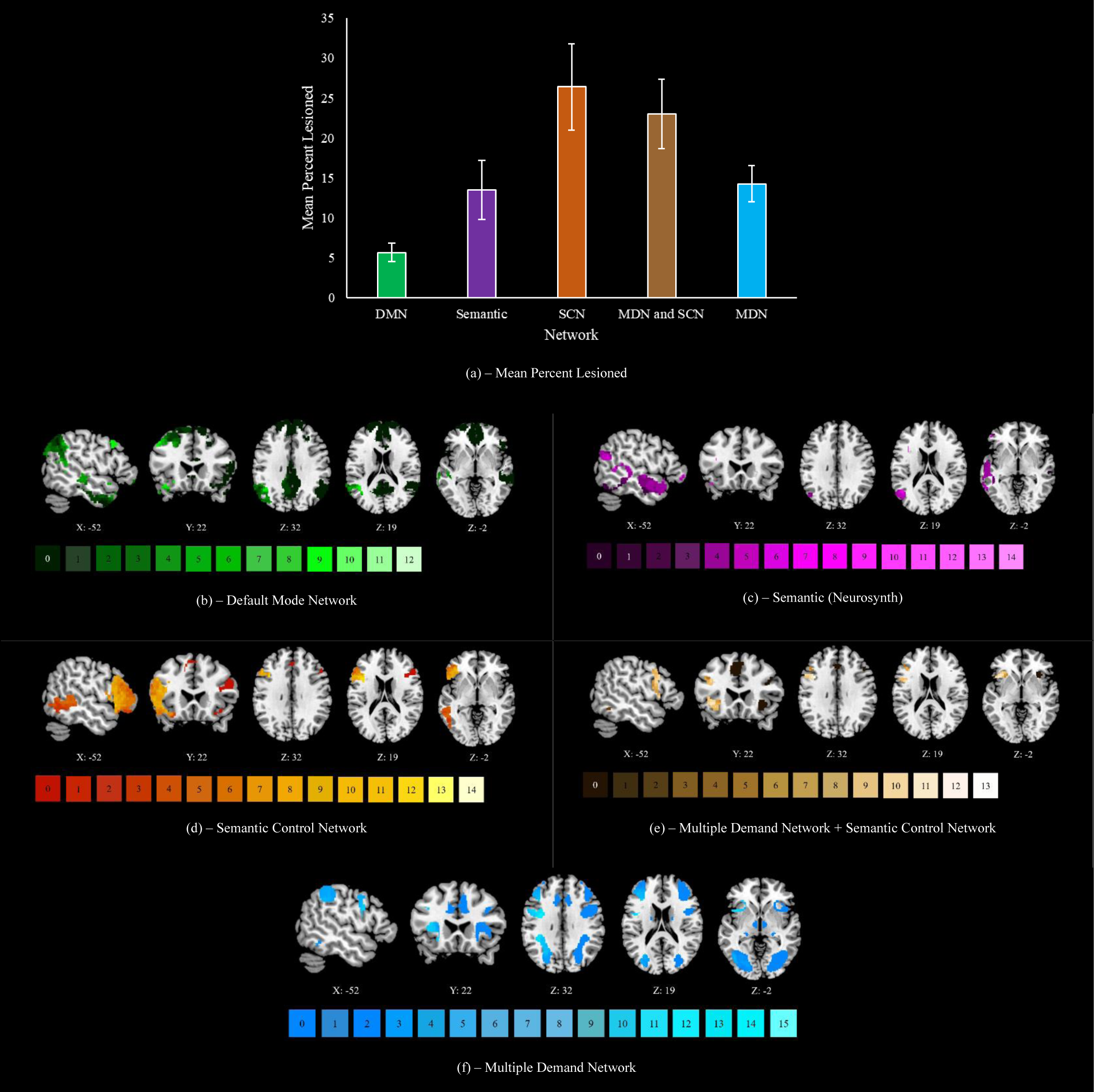
(a) The mean percentage of each network of interest (confined to the left hemisphere) overlapping with patient lesion files. DMN = default mode network, SCN = semantic control network, MDN = multiple demand network. Networks are visualised for (b) the DMN, (c) core semantic regions, (d) the SCN, (e) areas common to the MDN and SCN, and (f) the MDN. Any right hemisphere aspects of each network are visualised but were never impacted by lesion. Keys under each map reflect the number of patients with lesion to a given voxel. N = 23. DESCRIPTIVE CAPTION FOR TEXT-TO-SPEECH: Figure 2a reflects the mean percentage lesioned for each network of interest. This peaks in the semantic control network at 26%, followed by areas shared between the multiple demand and semantic **control networks at 23%, core semantic regions and regions exclusive to the multiple demand network both at 14%, and the default mode network at 6%. Locations of most frequent damage are displayed for each network in following sections.** **Figure 2b** **presents the default mode network, lesion peaks in the angular gyrus and insular cortex at a threshold of 12 cases.** **Figure 2c** **presents core semantic regions, lesion peaks in the inferior frontal gyrus pars opercularis at a threshold of 14 cases.** **Figure 2d** **presents the semantic control network, lesion peaks in the inferior frontal gyrus pars opercularis at a threshold of 14 cases.** **Figure 2e** **presents regions shared by the semantic control and multiple demand networks, lesion peaks in the middle frontal and precentral gyri at 13 cases.** **Figure 2f** **presents the multiple demand network, lesion peaks in the precentral gyrus at a threshold of 15 cases.**

#### 3.4.2. Spreading SCN disconnection

In order to understand typical damage to key semantic control regions in SA, we next identified SCN components most likely to be lesioned across the sample and looked for evidence of spreading structural and functional disconnection to SCN regions beyond the lesion site. The SCN map was split into its distinct clusters (see Section 2.7). Table 1 presents the percentage of each cluster lesioned for each patient, as well as a binary measure of whether this cluster showed evidence of structural or functional disconnection. Most cases had lesions in the left frontal cluster (16/23). All patients had structural and functional disconnection to this cluster, including when this cluster was not itself lesioned. A smaller number of cases had a lesion encompassing a left posterior temporal cluster (10/23) but again, all patients showed evidence of its structural and functional disconnection. A majority of cases showed evidence of structural (19/23) and functional disconnection (22/23) to the bilateral dmPFC cluster, despite only two cases showing direct lesion to this site. Four patients did not show direct lesion to any of the three left hemisphere SCN nodes. Even in these cases, diffuse connection was still frequently observed across the network. The two right hemisphere frontal clusters were spared in terms of lesion and structural disconnection but were both functionally disconnected in all but one patient.

**Table 1.** Lesion, structural disconnection (SDC), and functional disconnection (FDC) to semantic control network clusters.

| Patient | Left IFG, insula, OFC, & precentral gyrus |  |  | Left pMTG, pITG, & pFG |  |  | Bilateral dmPFC |  |  | Right IFG (pars orbitalis) |  |  | Right IFG (pars triangularis) |  |  |
| --- | --- | --- | --- | --- | --- | --- | --- | --- | --- | --- | --- | --- | --- | --- | --- |
|  | Lesion | SDC | FDC | Lesion | SDC | FDC | Lesion | SDC | FDC | Lesion | SDC | FDC | Lesion | SDC | FDC |
| P01 | ✗ | ✓ | ✓ | 46% | ✓ | ✓ | ✗ | ✗ | ✓ | ✗ | ✗ | ✓ | ✗ | ✗ | ✓ |
| P02 | 29% | ✓ | ✓ | 39% | ✓ | ✓ | ✗ | ✓ | ✓ | ✗ | ✗ | ✓ | ✗ | ✗ | ✓ |
| P03 | ✗ | ✓ | ✓ | ✗ | ✓ | ✓ | ✗ | ✗ | ✓ | ✗ | ✗ | ✓ | ✗ | ✗ | ✓ |
| P04 | 45% | ✓ | ✓ | 20% | ✓ | ✓ | ✗ | ✓ | ✓ | ✗ | ✗ | ✓ | ✗ | ✗ | ✓ |
| P05 | ✗ | ✓ | ✓ | 9% | ✓ | ✓ | ✗ | ✓ | ✓ | ✗ | ✗ | ✓ | ✗ | ✗ | ✓ |
| P06 | 48% | ✓ | ✓ | ✗ | ✓ | ✓ | ✗ | ✓ | ✓ | ✗ | ✗ | ✓ | ✗ | ✗ | ✓ |
| P07 | ✗ | ✓ | ✓ | 3% | ✓ | ✓ | ✗ | ✗ | ✓ | ✗ | ✗ | ✓ | ✗ | ✗ | ✓ |
| P08 | 80% | ✓ | ✓ | 25% | ✓ | ✓ | ✗ | ✓ | ✓ | ✗ | ✗ | ✓ | ✗ | ✗ | ✓ |
| P09 | 8% | ✓ | ✓ | ✗ | ✓ | ✓ | ✗ | ✓ | ✓ | ✗ | ✗ | ✓ | ✗ | ✗ | ✓ |
| P10 | 72% | ✓ | ✓ | 38% | ✓ | ✓ | ✗ | ✓ | ✓ | ✗ | ✗ | ✓ | ✗ | ✗ | ✓ |
| P11 | 48% | ✓ | ✓ | 1% | ✓ | ✓ | ✗ | ✓ | ✓ | ✗ | ✗ | ✓ | ✗ | ✗ | ✓ |
| P12 | 63% | ✓ | ✓ | ✗ | ✓ | ✓ | 3% | ✓ | ✓ | ✗ | ✗ | ✓ | ✗ | ✗ | ✓ |
| P13 | 19% | ✓ | ✓ | ✗ | ✓ | ✓ | ✗ | ✓ | ✓ | ✗ | ✗ | ✓ | ✗ | ✗ | ✓ |
| P14 | ✗ | ✓ | ✓ | ✗ | ✓ | ✓ | ✗ | ✓ | ✗ | ✗ | ✗ | ✗ | ✗ | ✗ | ✓ |
| P15 | 5% | ✓ | ✓ | ✗ | ✓ | ✓ | ✗ | ✓ | ✓ | ✗ | ✗ | ✓ | ✗ | ✗ | ✓ |
| P16 | 24% | ✓ | ✓ | 54% | ✓ | ✓ | ✗ | ✓ | ✓ | ✗ | ✗ | ✓ | ✗ | ✗ | ✓ |
| P17 | ✗ | ✓ | ✓ | ✗ | ✓ | ✓ | ✗ | ✓ | ✓ | ✗ | ✗ | ✓ | ✗ | ✗ | ✓ |
| P18 | 35% | ✓ | ✓ | ✗ | ✓ | ✓ | ✗ | ✓ | ✓ | ✗ | ✗ | ✓ | ✗ | ✗ | ✓ |
| P19 | 64% | ✓ | ✓ | ✗ | ✓ | ✓ | 4% | ✓ | ✓ | ✗ | ✗ | ✓ | ✗ | ✗ | ✓ |
| P20 | 20% | ✓ | ✓ | ✗ | ✓ | ✓ | ✗ | ✓ | ✓ | ✗ | ✗ | ✓ | ✗ | ✗ | ✓ |
| P21 | ✗ | ✓ | ✓ | ✗ | ✓ | ✓ | ✗ | ✗ | ✓ | ✗ | ✗ | ✓ | ✗ | ✗ | ✓ |
| P22 | 82% | ✓ | ✓ | 46% | ✓ | ✓ | ✗ | ✓ | ✓ | ✗ | ✗ | ✓ | ✗ | ✗ | ✓ |
| P23 | 93% | ✓ | ✓ | ✗ | ✓ | ✓ | ✗ | ✓ | ✓ | ✗ | ✗ | ✓ | ✗ | ✗ | ✓ |
| Group Mean | 34% | N/A | N/A | 9% | N/A | N/A | 0.3% | N/A | N/A | 0% | N/A | N/A | 0% | N/A | N/A |
| Patients Impacted | 16 | 23 | 23 | 10 | 23 | 23 | 2 | 19 | 22 | 0 | 0 | 22 | 0 | 0 | 23 |

Note: Semantic control network clusters taken from Jackson (2021). Lesion columns reflect the percentage of each cluster impacted by a given patient's lesion. In SDC and FDC columns, ticks reflect cases where there was any overlap with a cluster and patients' structural disconnection and functional disconnection maps, respectively. Crosses occur when a given cluster was not lesioned/disconnected. Patients Impacted reflects the number of patients for whom the given cluster was lesioned/disconnected. IFG = inferior frontal gyrus, OFC = orbitofrontal cortex, pMTG = posterior middle temporal gyrus, pITG = posterior inferior temporal gyrus, pFG = posterior fusiform gyrus, dmPFC = dorsomedial prefrontal cortex, SDC = structural disconnection, FDC = functional disconnection.

Overall, these results suggest maximum impact to areas implicated in semantic control in SA patients, with some damage to core semantic and domain-general control and relative sparing of the DMN. Patients reliably show evidence of spreading structural and functional disconnection to left hemisphere sites in the SCN that are not directly lesioned.

### 3.5. Symptom mapping

Next, we identified lesioned and disconnected voxels associated with poorer behavioural performance. Damage or disconnection was used to predict lower scores on the semantic cognition composite (comprising word and picture semantic associations and comprehension of ambiguous words) and the Brixton Spatial Anticipation Test, which probes non-verbal cognitive control, with results thresholded at t > 2.6. At this threshold, clusters associated with semantic and executive performance were mutually exclusive with no overlapping voxels within each analysis. We also examined the extent to which the clusters found in lesion-symptom mapping were implicated in our functional networks of interest, shown visual representations of which can be seen in Figure 2.

#### 3.5.1. Lesion-symptom mapping

Lesioned clusters associated with lower semantic cognition composite and Brixton scores can be seen in Figure 3a. Clusters implicated in semantic cognition showed overlap with the SCN, particularly in the anterior IFG, MFG, frontal pole, pMTG, and temporo-occipital part of the MTG. Overlap with the MDN was observed in the posterior IFG/inferior frontal sulcus, MFG, frontal pole, inferior LOC, intraparietal sulcus, and superior parietal lobule. Overlap with both the DMN and core semantic regions was observed in the frontal pole, pMTG, temporo-occipital part of the MTG, and temporo-occipital part of the ITG. Outside of these networks, clusters were observed in the precentral gyrus, postcentral gyrus, and occipital pole. Clusters associated with poorer performance on the Brixton test showed overlap with the SCN in the posterior IFG, MFG, SFG, and precentral gyrus. Overlap with the MDN was observed in intraparietal sulcus, SG and postcentral gyrus. Overlap with the DMN occurred in the SFG, while core semantic regions were not implicated. The percentage of each network of interest (restricted to the left hemisphere) implicated in these lesion-symptom maps is shown in Supplementary Figure 6.

**Figure 3.**
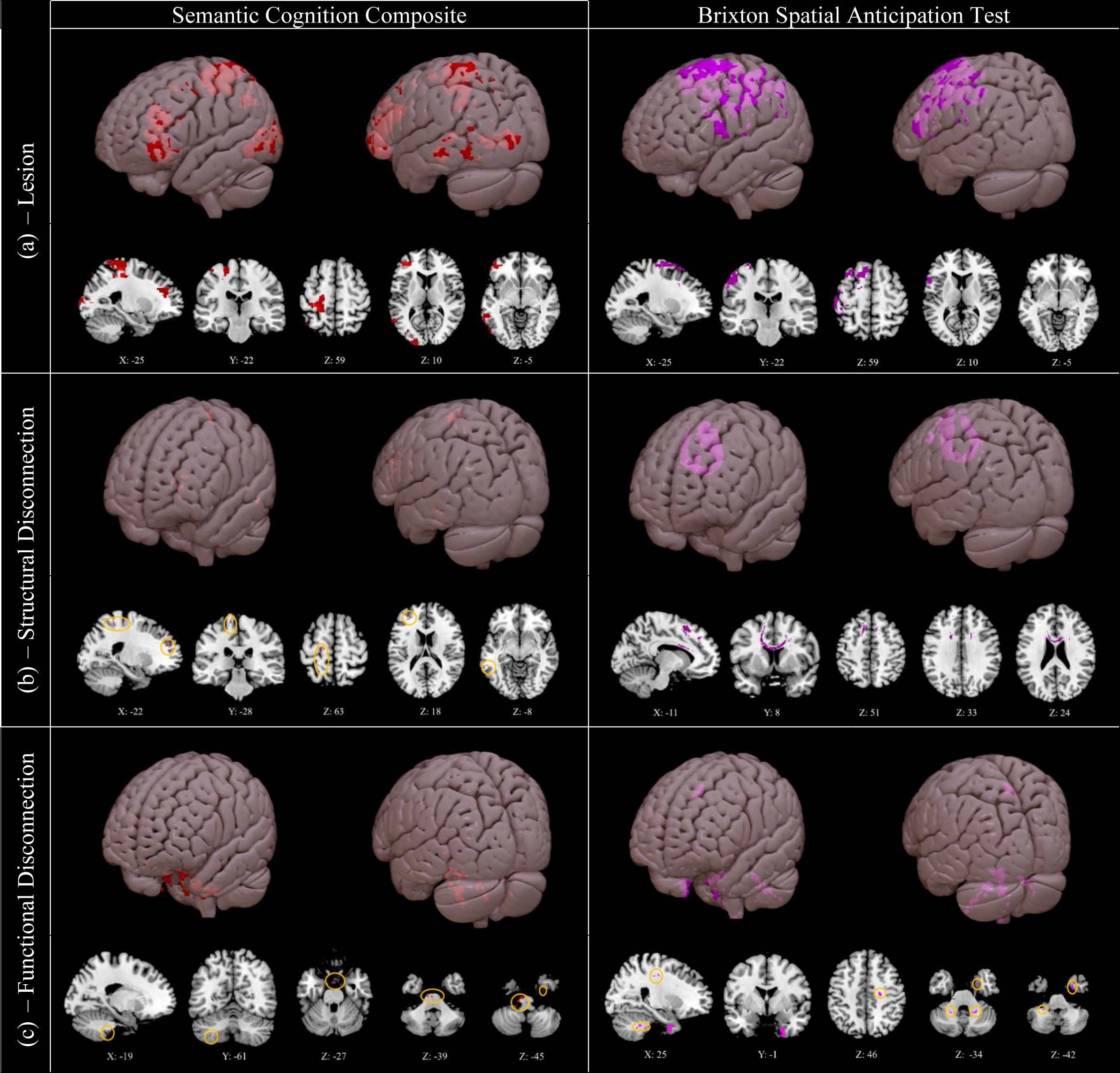
Clusters associated with lower semantic cognition composite scores (left) and lower scores on the Brixton Spatial Anticipation Test (right), for (a) lesion, (b) structural disconnection, and (c) functional disconnection data. Generated using non-parametric permutation tests in Randomise with threshold-free cluster enhancement. Highlighted voxels have a t-value of 2.6 or higher. Small clusters are highlighted in orange circles. 3D rendering generated in SurfIce. N = 20. DESCRIPTIVE CAPTION FOR TEXT-TO-SPEECH: Figure 3 presents clusters identified in symptom mapping as associated with poorer performance. This is done for both the semantic composite score and Brixton Spatial Anticipation Test, for the measures of lesion, structural disconnection, and functional disconnection. Lesioned clusters associated with poorer semantic and Brixton performance are large and left lateralised, and reflect the regions listed in section 3.5.1. Structurally disconnected clusters are small and left lateralised for semantic cognition, reflecting regions listed in section 3.5.2., while for poorer Brixton performance one large cluster is observed across the corpus callosum. Functionally disconnected clusters are small for both measures, and reflect the regions listed in section 3.5.3.

#### 3.5.2. Structural disconnection-symptom mapping

The results of the structural disconnection analysis can be seen in Figure 3b.

Structurally disconnected clusters associated with lower semantic cognition composite scores were minimal but were confined to the left hemisphere in regions including the frontal pole, precentral and postcentral gyri, pMTG, and occipital pole. These clusters were proximal to the regions identified in the lesion analysis but were not large enough to implicate specific white matter tracts and may not have clinical significance. There were no structurally disconnected clusters associated with poor semantic cognition in the right hemisphere. In contrast, poorer Brixton performance was associated with a pattern of interhemispheric structural disconnection across the corpus callosum, consistent with a role of interhemispheric connectivity in executive control in this visuo-spatial task.

#### 3.5.3. Functional disconnection-symptom mapping

The results of the functional disconnection analysis are shown in Figure 3c. Lower semantic cognition composite scores and poorer Brixton performance were both predicted by small functionally disconnected clusters in the brainstem, bilateral cerebellum, right parahippocampal gyrus, and right temporal fusiform cortex. Poorer Brixton performance was also associated with functional disconnection of the right temporal pole and of white matter proximal to the right precentral gyrus. In both cases, these clusters were sparse and too small to implicate functional networks; consequently, they may not have clinical significance.

In summary, left hemisphere lesion sites associated with poorer semantic cognition are mutually exclusive from but adjacent to those implicated in executive function (in line with previous studies showing that the semantic control network lies between DMN and MDN regions on the cortical surface; Davey et al., 2016; Wang et al., 2020). In contrast, the substrates for structural disconnection are more divergent across semantic and executive tasks, since the small clusters associated with poor semantic cognition are left-lateralised, while executive dysfunction is associated with cross-hemispheric disconnection. Clusters identified from functional disconnection symptom mapping were small and did little to distinguish between semantic and executive performance.

## 4. Discussion

This study characterised lesion location, structural disconnection, and functional disconnection in semantic aphasia (SA) patients who have impaired semantic control following left hemisphere stroke. Lesions were most common within the semantic control network (SCN), and areas commonly responsive to semantic control and executive function (the multiple demand network (MDN); Duncan, 2010). Lesions were heterogenous as is typical of this population (e.g., Chapman et al., 2020; Hallam et al., 2018), with many but not all cases having frontal damage. Measures of structural and functional disconnection, anticipated from lesion location, showed extensive effects spreading beyond the lesion, frequently affecting structurally-intact SCN regions. This may account for the similarity of control deficits seen in SA patients despite divergent lesion profiles.

When mapping behavioural performance to lesion site, more severe deficits of semantic cognition and non-semantic cognitive control were associated with lesions in adjacent areas of the left frontoparietal cortex – consistent with studies from healthy participants suggesting that semantic and domain-general executive regions are supported by adjacent left hemisphere regions (Davey et al., 2016; Gao et al., 2021). Associations between functional disconnection and cognitive performance were limited and did not identify distinct patterns for semantic and executive deficits. In contrast, semantic and executive deficits had different associations with structural disconnection. While insight into semantic performance from structural disconnection was limited, with small frontal, parietal, and posterior temporal clusters in the left hemisphere, proximal to lesioned substrates, being highlighted, executive dysfunction was predicted by interhemispheric structural disconnection across the corpus callosum. These different patterns are consistent with previous studies showing that the SCN is largely left-lateralised, while domain-general executive function is supported by the bilateral MDN (Fedorenko et al., 2013; Gonzalez Alam et al., 2019). In this way, we can explain why similar left hemisphere clusters are associated with poor semantic cognition and cognitive control and provide an account of the hemispheric differences in structural disconnection associated with these deficits.

Previous descriptions of the SCN have highlighted left IFG and pMTG as key structures (Jackson, 2021; Jefferies, 2013; Jefferies et al., 2019; Lambon Ralph et al., 2017; Noonan et al., 2013). Patients in the current sample frequently presented with direct damage to these SCN sites, and invariably had some degree of structural and functional disconnection within this network. This pattern was observed even in four patients who showed no direct SCN damage; in these cases, semantic impairment may be attributable to SCN disconnection (although direct effects of other areas of lesion might also be relevant in regions involved in semantic cognition yet omitted from prior meta-analyses of semantic control). This damage and disconnection of SCN regions in SA is consistent with neuroimaging evidence implicating left IFG and pMTG in semantic control (Jackson, 2021; Noonan et al., 2013).

Topographically, the SCN lies between the DMN and MDN in the left hemisphere (Davey et al., 2016; Wang et al., 2020), and this network has been proposed as a functional nexus between core semantic and domain-general control regions (Davey et al., 2016). Accordingly, the current results revealed that adjacent but distinct frontoparietal lesions resulted in poorer semantic cognition and executive function, consistent with prior evidence of adjacent neural substrates for these functions (Wang et al., 2018). This may account for why some SA patients present with executive dysfunction while others do not (Thompson et al., 2018).

Indeed, 14 of the current 23 patients presented with some degree of executive dysfunction, with performance on the Brixton Spatial Anticipation Test positively correlating with semantic performance. It may be that impaired semantic control does not necessitate executive impairment (Chapman et al., 2020), but that the proximity of these substrates means that these functions are frequently impaired together. Frequent damage to substrates underlying semantic or domain-general control or both in SA may give rise to broad deficits in constraining internal aspects of cognition (Souter et al., 2021; Stampacchia et al., 2018), reflected in heightened susceptibility to external cues and miscues in semantic retrieval (Jefferies et al., 2008; Noonan et al., 2010), along with strong effects of distractors in semantic decision-making (Corbett et al., 2011; Noonan et al., 2010).

While the lesioned substrates associated with semantic cognition included SCN regions as expected, areas outside this network were also implicated, including the LOC and postcentral gyrus. It is worth noting that the semantic composite score derived here reflects general semantic cognition (albeit performance on highly demanding heteromodal tasks) rather than semantic control specifically, and this may account for the inclusion of regions not typically associated with controlled processing. Evidence has suggested the involvement of the LOC in processing visual features of concepts (Coutanche & Thompson-Schill, 2015), while the postcentral gyrus has been implicated in sensorimotor representation (Kropf et al., 2019). Furthermore, the LOC has been shown to decode category- and task-related semantic information (Wang et al., 2021). Similarly, the postcentral gyrus has been implicated in the performance of divergent thinking tasks, which require the generation of alternative uses of objects (Cogdell-Brooke et al., 2020). Such a process is comparable to semantic control in requiring focus on subordinate conceptual information. Finally, the LOC and postcentral gyrus are adjacent to the pMTG and dorsal AG, respectively; two nodes that have been implicated in semantic control (Hodgson et al., 2021; Noonan et al., 2013). Ultimately, while the functional contribution of the LOC and postcentral gyrus are not fully understood, these regions may support conceptual retrieval by either representing features of concepts or by supporting semantic decision-making.

In previous studies, several white matter tracts have been associated with semantic cognition, including the IFOF, ATR, UF, and ILF (Almairac et al., 2015; Han et al., 2013; Sierpowska et al., 2019). These tracts were reliably disconnected across the sample, as were others including the SLF, AF, and FAT. The SLF and AF have both been implicated in language processing (Catani & Mesulam, 2008; Han et al., 2013), and the FAT in both language and executive function (Dick et al., 2019). However, the clusters we identified in the semantic cognition structural disconnection symptom mapping were too small to implicate any specific tracts. This may in part be because the pattern of structural disconnection across the sample was diffuse and widespread; multiple tracts underpin semantic cognition and damage to any one of them may be sufficient to produce an impairment of semantic cognition. This analysis may have also been hampered by insufficient statistical power, due to our relatively small sample size. Despite this, our findings are consistent with emerging accounts of the lateralisation of control functions.

Clusters associated with demanding semantic tasks were small and left-lateralised, intersecting with the SCN. Contrasts of hard over easy semantic judgements in studies of healthy participants also reveal a highly left-lateralised network (Gonzalez Alam et al., 2019; Jackson, 2021; Noonan et al., 2013). In contrast, the ventral ATL’s role in long-term semantic representation is highly bilateral (Gonzalez Alam et al., 2019; Rice et al., 2015).

Conversely, executive performance on a demanding visual-spatial task was predicted by interhemispheric structural disconnection across the corpus callosum, indicating that this task may be supported by a more bilateral network, in line with prior work (e.g., Bodini et al., 2013; Johnson et al., 2017; Schulte & Müller-Oehring, 2010). Neuroimaging studies show that bilateral activation underlies executive function (Camilleri et al., 2018; Fedorenko et al., 2013) and (especially in visual-spatial tasks), the most robust responses are often right-lateralised (Dick et al., 2019; Gonzalez Alam et al., 2018). Moreover, a recent study found that control networks show different patterns of connectivity across the hemispheres, with left-lateralised control regions showing stronger connectivity with heteromodal regions of the DMN (Gonzalez Alam et al., 2021). In this way, the left lateralised nature of the semantic control network may be adaptive, allowing control regions to separate from visuospatial responses when internal aspects of cognition are constrained.

Prior studies investigating the predictive value of indirect measures of structural disconnection (inferred from lesion location, as in the current study) have yielded inconsistent results. Salvalaggio et al. (2020) provided evidence that structural disconnection is comparable to lesion location in its ability to predict post-stroke deficits across domains. Halai et al. (2020) directly measured structural disconnection using diffusion weighted imaging data, and conversely found no added benefit relative to direct observations of abnormal tissue when predicting post-stroke aphasia, citing tight coupling between lesion and disconnection as a possible cause. This is consistent with our observation that the small clusters of structural disconnection associated with deficits in semantic cognition largely followed lesioned substrates, although a different pattern was found for executive dysfunction. Hope et al. (2018) similarly found limited predictive value of indirectly measured structural disconnection beyond lesion location for predicting aphasia severity, while other studies have suggested unique contributions from such measures (Del Gaizo et al., 2017; Kristinsson et al., 2021). It may be that structural disconnection is unlikely to explain unique variation in behavioural performance beyond lesion location when patients have been selected to show particular deficits associated with areas of cortex that are lesioned, as was the case for semantic deficits in our sample. However, since stroke lesions are typically unilateral, structural disconnection may be more likely to explain unique variance in other domains associated with bilateral networks, including executive dysfunction. This is consistent with prior evidence demonstrating the utility of structural disconnection in reflecting inter-hemispheric networks (Thiebaut de Schotten et al., 2020).

Clusters highlighted by functional disconnection were limited in the current study, but included the bilateral cerebellum, brainstem, and right ventral temporal regions across both semantic and executive measures. Unlike structural disconnection, patterns of functional disconnection did not distinguish between our two behavioural domains. While the cerebellum is reliably activated in neuroimaging studies (Buckner, 2013; Stoodley & Schmahmann, 2009), other sites such as medial temporal cortex have relatively low signal-to-noise (e.g., Olman et al., 2009) and the validity of these observed clusters is therefore unclear. The appropriateness of using indirect estimates of functional disconnection to predict behavioural deficits, and the methods that can be used to determine these estimates, also remain contentious (Boes, 2021; Bowren et al., 2022; Salvalaggio et al., 2021a; Umarova & Thomalla, 2020). Some studies have found that indirect measures of functional disconnection have relatively poor utility for predicting behavioural deficits post-stroke compared with measures of structural disconnection (Salvalaggio et al., 2020); our findings do not challenge this view, although other studies report more success (Cohen et al. 2021; Ferguson et al., 2019; Padmanabhan et al., 2019). Discrepancies between studies may reflect differences in data pre-processing or the methods used to estimate the accuracy of behavioural predictions (Salvalaggio et al., 2021b). Therefore, our lack of meaningful results might reflect methodological limitations, even though we employed recent recommendations for estimating functional disconnection, using PCA to identify connectivity patterns within each patients’ lesion, rather than seeding the entire lesioned region. This approach has been reported to increase functional specificity and support behavioural prediction (Pini et al., 2021).

There are some important limitations of this study. First, the current sample size was relatively small, due to practical barriers in recruiting, testing, and scanning patients who fit the criteria for semantic aphasia. Secondly, as data were collated across three samples of SA patients, there were a limited number of tests on which all patients had been assessed. Our analysis of executive function was confined to the Brixton Spatial Anticipation Test. While performance on Raven’s Coloured Progressive Matrices (Raven, 1962) was available for each patient, scores across these two tests did not meet the necessary assumptions of PCA, and so a composite score was not derived. While the Brixton test is a sensitive measure of executive impairment (van den Berg et al., 2009), the use of a broader set of tests would be helpful in establishing the relationship between damage and general deficits of cognitive control. Third, measures used to examine impaired semantic cognition were not specific to semantic control.

Prior investigations have consistently concluded that SA patients present with preserved semantic storage and impaired controlled retrieval (e.g., Jefferies & Lambon Ralph, 2006; but see Chapman et al., 2020 for an alternative view). However, in the absence of a comparison group we cannot rule out the possibility that aspects of our results reflect impaired semantic processing more generally. A comparison of disconnection in SA and SD may provide additional insights into how these conditions differentially impact networks associated with controlled retrieval and core semantic representation. SD patients show disrupted functional connectivity between the left anterior hippocampus, ATL/insula and MTG (Chen et al., 2017), but a group comparison of SD and SA cases is not yet available. A study directly comparing structural connectivity in SD and SA cases found semantic performance is predicted by fractional anisotropy of the left medial temporal network in SD, as opposed to left fronto-subcortical and fronto-temporal/occipital networks in SA (Ding et al., 2020); however, network differences linked to conceptual representation and control may be confounded by differences in aetiology between these groups, since stroke frequently damages white-matter tracks, while the knife-edge atrophy in SD primarily affects grey matter. Indeed, Andreottia et al. (2017) provide evidence that that structural network disruption in SD is localised to regions directly affected by atrophy, suggesting that symptom mapping from such a measure may provide limited additional insight. A final limitation is that our measures of functional and structural disconnection were indirect. Direct measures of resting state functional connectivity have been shown to predict post-stroke symptoms to a greater extent than indirectly measured functional disconnection (Salvalaggio et al., 2020).

### 4.1. Conclusion

We assessed patterns of spreading structural and functional disconnection in left hemisphere stroke patients with semantic aphasia. These results highlight damage to the SCN in this group, both as a direct result of lesions and following spreading disconnection. We show that semantic and domain-general executive control are supported by adjacent substrates in the left hemisphere and yet associated with distinct patterns of structural disconnection that are left-lateralised and bilateral, respectively.

## Acknowledgements

We thank Zhiyao Gao for his assistance in the procurement and interpretation of functional network maps, and Lucy Cogdell-Brooke for her guidance in manual lesion tracing. EJ was funded by a European Research Council Consolidator grant (FLEXSEM – 771863). MTS was also funded by a European Research Council Consolidator grant (DISCONNECTOME, Grant agreement No. 818521). AH was funded by the Rosetrees Trust (A1699) and European Research Council (GAP: 670428 – BRAIN2MIND_NEUROCOMP to MLR).

## Statements & Declarations

### Author Contributions (CRediT authorship contribution statement)

Nicholas E. Souter: Conceptualization, Methodology, Formal analysis, Investigation, Data Curation, Writing – Original Draft, Writing – Review & Editing, Visualization, Project administration. Xiuyi Wang: Formal analysis, Writing – Review & Editing. Hannah Thompson: Investigation, Resources, Writing – Review & Editing. Katya Krieger-Redwood: Formal analysis, Writing – Review & Editing. Michel Thiebaut de Schotten: Conceptualization, Methodology, Software, Formal analysis, Resources, Writing – Review & Editing. Ajay D. Halai: Resources, Writing – Review & Editing. Matthew A. Lambon Ralph: Resources, Writing – Review & Editing. Elizabeth Jefferies: Conceptualization, Methodology, Investigation, Writing – Review & Editing, Supervision, Funding Acquisition.

### Data Availability

Data for this project are available on the Open Science Framework (https://osf.io/6psqj/). Unthresholded group-level NIFTI files corresponding to the results presented here can be seen on Neurovault (https://neurovault.org/collections/10333/). Raw neuroimaging data for individual patients is not published here due to insufficient consent.

### Ethics approval

This study was performed in line with the principles of the Declaration of Helsinki. Approval was granted by the York Neuroimaging Centre at the University of York (date: 24/10/2019, project ID: P1363).

### Consent to participate

Informed consent was obtained from all individual participants included in the study.

### Consent to publish

The authors affirm that human research participants provided informed consent for publication of all data included within this manuscript, supplementary materials, and open data repositories.

## Supplementary Materials Descriptive Captions

Supplementary Figure 1 DESCRIPTIVE CAPTION FOR TEXT-TO-SPEECH: Supplementary Figure 1 provides replication of the structural disconnection symptom mapping from the paper, but at alternative probability thresholds. This includes 40% in Supplementary Figure 1a, 50% in Supplementary Figure 1b, which is the default threshold used in the paper, and 60% in Supplementary Figure 1c. In each case, symptom mapping is presented for patients’ Semantic Cognition Composite Score on the left, and performance on the Brixton Spatial Anticipation Test on the right. Results are consistent at each threshold. For the Semantic Cognition Composite Score, this includes very small voxels located in the left the frontal pole, precentral and postcentral gyri, pMTG, and occipital pole. The only conceptual deviation from the default threshold is a single voxel highlighted in the right precentral gyrus at a threshold of 40%. Clusters are too small to provide clinical significance. For the Brixton Spatial Anticipation Test, at each threshold performance is predicted by structural disconnection across the corpus callosum.

Supplementary Figure 2 DESCRIPTIVE CAPTION FOR TEXT-TO-SPEECH: Supplementary Figure 2a reflects the mean percentage functionally disconnected for each network of interest. This peaks in the semantic control network at 85%, followed by areas shared between the multiple demand and semantic control networks at 81%, core semantic regions at 80%, regions exclusive to the multiple demand network both at 75%, and the default mode network at 44%. Locations of most frequent damage are displayed for each network in following sections. Supplementary Figure 2b presents the default mode network, functional disconnection peaks in the right inferior frontal gyrus (pars triangularis), left frontal orbital cortex, left temporooccipital part of the middle temporal gyrus, and left posterior supramarginal gyrus. Supplementary Figure 2c presents core semantic regions, functional disconnection peaks in the left temporooccipital part of the inferior temporal gyrus, left temporooccipital part of the middle temporal gyrus, and left inferior lateral occipital cortex. Supplementary Figure 2d presents the semantic control network, functional disconnection peaks in the bilateral inferior frontal gyrus (pars opercularis), and left temporooccipital part of the inferior temporal gyrus. Figure 2e presents region s shared by the semantic control and multiple demand networks, functional disconnection peaks in the left precentral gyrus, left inferior frontal gyrus (pars triangularis), and left temporooccipital part of the inferior temporal gyrus. Supplementary Figure 2f presents the multiple demand network, functional disconnection peaks in the bilateral precentral gyrus, right inferior frontal gyrus (pars opercularis) and left temporooccipital part of the inferior temporal gyrus.

Supplementary Figure 3 DESCRIPTIVE CAPTION FOR TEXT-TO-SPEECH: Supplementary Figure 3 presents clusters identified in symptom mapping as associated with relatively better performance. This is done for both the semantic composite score and Brixton Spatial Anticipation Test, for the measures of lesion, structural disconnection, and functional disconnection. Lesioned clusters associated with better semantic cognition are left fronto-parietal, and implicate the precentral, postcentral, and superior frontal gyri. Lesioned clusters associated with better Brixton performance include the frontal pole, planum temporale, angular gyrus, and postcentral gyrus. Structurally disconnected clusters associated with better semantic cognition include a small bilateral group of voxels in the parietal cortex which do not implicate specific regions or tracts, as well as a cluster in the left Heschl’s gyrus. Structurally disconnected clusters associated with better Brixton performance are similarly sparse, but implicate the putamen, occipital pole, superior parietal lobule and precentral gyrus. Functionally disconnected clusters associated with better semantic cognition include several sparse voxels, including in the brain stem, right temporal pole, and in white matter proximal to the right precentral gyrus. No clusters met the threshold of t > 2.6 for functionally disconnected clusters associated with better Brixton performance.

Supplementary Figure 4 DESCRIPTIVE CAPTION FOR TEXT-TO-SPEECH: Supplementary Figure 4 presents unthresholded overlap maps for the measures of lesion, structural disconnection, and functional disconnection. Supplementary Figure 4a presents lesion overlap. Lesions are confined to the left hemisphere and subsume much of the cortex, affecting each lobe, peaking in the precentral and middle frontal gyri. Maximum overlap is 16 cases. Supplementary Figure 4b presents structural disconnection overlap. Structural disconnection is largely left lateralised but with some spreading to the right hemisphere. Most left hemisphere white matter is implicated here, but this peaks in the left superior longitudinal fasciculus and inferior fronto-occipital fasciculus. Maximum overlap is all 23 cases. Supplementary Figure 4c presents functional disconnection overlap. Functional disconnection is bilateral and extensive, subsuming almost the entirety of the brain. This disconnection peaks in the left temporooccipital part of the inferior temporal gyrus, and right inferior frontal gyrus (pars opercularis). Maximum overlap is all 23 cases.

Supplementary Figure 5 DESCRIPTIVE CAPTION FOR TEXT-TO-SPEECH: Supplementary Figure 5a provides the mean probability that each tract of interest, as listed in section 2.5. of the manuscript, is disconnected across the sample. The mean probability of disconnection is highest in the corpus callosum at .99, followed by the superior longitudinal fasciculus 2 at .96, the arcuate fasciculus long at .95, and the superior longitudinal fasciculus 3 at .94. The lowest probability of disconnection is in the uncinate fasciculus at .61. Supplementary Figure 5b provides the mean estimated proportion disconnected for each tract. This is highest in the frontal inferior longitudinal fasciculus at .37, followed by the superior longitudinal fasciculus 3 at .32, and the anterior arcuate fasciculus at .30. The lowest estimated mean proportion disconnected is in the corpus callosum, at .05. Supplementary Figure 5c displays a visualisation of the structural disconnection overlap map for the sample, thresholded at 19 cases. Structural disconnection is left lateralised at this threshold and shows maximal overlap with the superior longitudinal fasciculus and inferior fronto-occipital fasciculus. Supplementary Figure 5d through Supplementary Figure 5k display visualisations of, respectively, the anterior thalamic radiation, uncinate fasciculus, inferior longitudinal fasciculus, frontal aslant tract, arcuate fasciculus, superior longitudinal fasciculus, inferior fronto-occipital fasciculus, and corpus callosum.

Supplementary Figure 6 DESCRIPTIVE CAPTION FOR TEXT-TO-SPEECH: Supplementary Figure 6a presents a bar graph of the percentage of left hemisphere voxels in each network of interest which are implicated in the group-level lesion symptom mapping output for the Semantic Cognition Composite Score. This peaks at 6.2% for the semantic control network, followed by 3.0% for the multiple demand network, 2.4% for core semantic regions, 0.6% for the default mode network, and 0.1% for areas shared by the semantic control and multiple demand networks. Supplementary Figure 6b provides the same graph for the Brixton Spatial Anticipation Test symptom mapping output. In this case, numbers peak in the semantic control network at 4.3%, followed by the default mode network at 3.6%, areas shared by the semantic control and multiple demand networks at 2.4%, and the multiple demand network at 2.3%, with 0% of core semantic regions implicated.

## Footnotes

1 vATL is typically spared in SA, since the anterior temporal cortical artery branches below the main trifurcation of the middle cerebral artery and because this watershed region has a dual blood supply from the middle and posterior cerebral arteries (Borden, 2006; Conn, 2003).

2 Age of leaving education missing for one participant (P19).

3 This measure was selected as it is a sensitive measure of executive impairment and was widely-available in our sample.

4 These anatomical regions were thresholded at a value of 30 such that they were mutually exclusive.

5 Numbers in this section reflect the percentage of the sample showing functional disconnection to the respective region.

6 Note that these average percentages will be impacted by differences in the relative size of each network. The average number of voxels lesioned over the total size of the respective number of voxels in each network is: DMN: 772/13,618, Semantic: 479/3,549, SCN: 934/3,538, MDN & SCN: 409/1,777, MDN: 1,817/12,731.

