## Supplementary Materials for "Mapping lesion, structural disconnection, and functional disconnection to symptoms in semantic aphasia"

### Background Neuropsychology

Patients completed a series of background tests probing language, memory, and executive function. Each individual patients' performance on these tests can be seen in Supplementary Table 1. Of the 15 patients tested, seven showed evidence of impaired word repetition using a subtest from the Psycholinguistic Assessments of Language Processing in Aphasia battery (PALPA; Kay et al., 1992). Of the 17 patients tested, 15 were impaired for category fluency (eight categories), while 16 were impaired for letter fluency (F, A, S). Sixteen of 22 patients presented with impaired forward digit span, while 14 of 19 presented with impaired backward digit span (Wechsler Memory Scale III; Wechsler, 1997). Eight patients presented with impairments in visuospatial processing, as measured by subtests of the Visual Object and Space Processing Battery (VOSP; Warrington & James, 1991). Patients also completed several tests of executive function, including a subtest of the Test of Everyday Attention (Robertson et al., 1994), Raven's Coloured Progressive Matrices (Raven, 1962), and the Brixton Spatial Anticipation Test (Burgess & Shallice, 1997). All patients completed at least one of these tests, with fourteen showing some evidence of impairment.

As a measure of core semantic ability, participants completed the Cambridge Semantic Battery (Bozeat et al., 2000). Each individual patient's performance on these tests can be seen in Supplementary Table 2. Of the 20 tested, 16 patients were impaired on the Picture Naming task [Mean (SD) = 54.6% (37.4)], in which they were required to verbally provide the name for a series of black and white line drawings. Though not a part of the Cambridge Semantic Battery, providing phonemic cues as to the correct target label improved all patients' performance to ceiling or near-ceiling level [Mean (SD) = 75.3% (41.5)]. Of the 21 tested, 12 patients showed impaired performance on Word-Picture Matching [Mean (SD) = 91.5% (9.4)], in which they were required to match one of ten possible line drawings to a verbally provided probe word. The Camel and Cactus Test (CCT) was used a measure of ability to make thematic associations, requiring matching a probe word/picture to one of four possible targets. 21 patients completed the full version of these tasks, while the remaining two (P22 and P23) completed shortened versions. Of those who completed the full task, 18 patients were impaired on the word version of the CCT [Mean (SD) = 74.1% (18.3)], while 12 were impaired on the picture version [Mean (SD) = 74.5% (21.8)]. The two patients who completed the short versions of the CCT were impaired on both the word and pictures versions.

The ambiguity task (Noonan et al., 2010) required patients to make thematic associations between a probe word and one of three possible targets. Each probe word was a homonym with a dominant (e.g., PEN – PENCIL) and subordinate (e.g., PEN – PIG) association. The latter is believed to tax semantic control more than the former, due to the need to flexibly retrieve non-dominant semantic information (Thompson et al., 2017). Probe words were either presented with no cue, with a contextual cue alluding to the correct target meaning of the word (e.g., PEN – PIG: "the labourers cleaned out the pen"), or with a miscue, alluding to the incorrect interpretation (e.g., PEN – PIG: "he signed his name with a fountain pen"). Twenty-one patients completed the no cue version of the task, with 14 also completing the cue and miscue versions. In the no cue condition, patients performed better for dominant [Mean (SD) = 79.0% (13.8)] than subordinate trials [Mean (SD) = 53.5% (15.1)]. Relative to no cue, cued trials improved performance on subordinate [Mean (SD) = 71.9% (15.6)] but not dominant trials [Mean (SD) = 77.4% (14.6)]. Miscued trials considerably impaired accuracy on dominant [Mean (SD) = 61.2% (21.3)], and somewhat on subordinate trials [Mean (SD) = 45.0% (19.9)]. Contextual cues therefore improved accuracy on the most difficult trials, while contextual miscues impaired performance on the easiest trials.

The synonym judgement task (Samson et al., 2007) required participants to match a probe word to a possible synonym, presented alongside two foils. In each trial, one of these foils acted as either a strong (e.g., probe: DESERT, target: WILDERNESS, distractor: SAND) or weak (e.g., probe: HAZARD, target: DANGER, distractor: LIGHT) thematic distractor. Strong thematic distractors should impair performance to a greater extent than weak distractors, as SA patients are strongly influenced by irrelevant but competing information (Jefferies, 2013). Sixteen patients were tested on this measure. Overall, the sample performed better on weak distractor trials [Mean (SD) = 69.3% (13.4)] than strong distractor trials [Mean (SD) = 49.6% (16.6)]. All but one patient (P13) showed this expected pattern.

The object use task (Corbett et al., 2011) provides a non-verbal measure of semantic control. Herein, patients are required to identify the appropriate object, of six possible options, to perform a given action (e.g., "Crack a nut"). The target objects could be either be 'canonical' such that they are typically used to complete this action (e.g., NUT CRACKER), or an 'alternative' object which could be used to complete the action if necessary (e.g., HAMMER). Alternative trials should require greater semantic control as they require access to non-dominant information about the target object, and inhibition of dominant information (e.g., that hammers are typically used in construction). Twenty patients were tested on this measure. Overall, the sample performed better on canonical [Mean (SD) = 92.7% (7.5)] than on alternative trials [Mean (SD) = 59.5% (19.7)]. This was true for all 20 patients.

*Supplementary Table 1. Patient performance on background neuropsychological testing.*

| | Language | | | Verbal working memory | | Executive | | | | Visual Object and Space Processing battery | | | |
|---|---|---|---|---|---|---|---|---|---|---|---|---|---|
| | PALA 9 Word repetition | Category Fluency | Letter Fluency | Forwards digit span | Backwards digit span | Brixton | Ravens | TEA without distraction | TEA with distraction | Dot counting | Position discrimination | Number location | Cube analysis |
| Max | 80 | - | - | 8 | 7 | 54 | 36 | 7 | 10 | 10 | 20 | 10 | 10 |
| Cut-off | 73 | 62 | 18 | 5.54 | 3.66 | 28 | 28 [a] | 4.2 | 2.6 | 8 | 18 | 7 | 6 |
| Mean | 62.2 | 32.3 | 7.8 | 3.6 | 1.8 | 23.7 | 26.0 | 5.0 | 4.1 | 8.7 | 18.4 | 8.3 | 6.8 |
| P01 | **68.8** | **49** | **14** | 6 | **2** | 28 | 20 | 7 | 9 | 8 | 19 | 9 | **4** |
| P02 | **64** | **18** | **0** | **4** | 2 | **7** | **12** | 6 | 3 | 10 | 18 | 9 | **3** |
| P03 | 75.2 | **24** | 19 | 8 | 4 | 28 | 31 | 5 | 9 | NT | NT | NT | NT |
| P04 | 76.8 | **11** | **8** | **4** | **1** | **14** | **6** | **2** | 3 | **6** | **16** | 8 | **4** |
| P05 | 80 | **25** | **14** | 6 | **3** | **11** | **13** | 7 | 9 | **3** | **15** | **2** | **4** |
| P06 | **64.8** | **25** | **5** | **3** | 2 | 34 | 26 | **3** | **2** | 10 | 20 | 10 | **5** |
| P07 | NT | **61** | **13** | **5** | 3 | 37 | 29 | 5 | 6 | NT | NT | NT | NT |
| P08 | NT | NT | NT | **0** | **0** | 34 | **24** | 5 | **1** | 8 | 20 | 8 | 9 |
| P09 | 75 | **26** | **2** | 5 | **2** | **26** | **24** | 5 | **1** | 9 | 19 | 10 | **4** |
| P10 | **42** | **15** | **2** | **1** | **0** | **18** | 31 | 6 | **1** | 10 | **15** | **5** | **4** |
| P11 | **1** | NT | NT | **2** | NT | 31 | 34 | 7 | 7 | 8 | 19 | 10 | 10 |
| P12 | **71** | **26** | **2** | **4** | **2** | **7** | **27** | 5 | 3 | 10 | 20 | 10 | 9 |
| P13 | **7** | 69 | **12** | 6 | 4 | 39 | 33 | 5 | 3 | 10 | 20 | 8 | 8 |
| P14 | 74 | 80 | **16** | **4** | **2** | 31 | **21** | 5 | **2** | 10 | 20 | **5** | 10 |
| P15 | NT | NT | NT | **0** | **0** | **21** | 31 | **2** | **1** | **7** | 19 | 8 | 8 |
| P16 | NT | **4** | **3** | **3** | NT | NT | 31 | 7 | 3 | NT | NT | NT | NT |
| P17 | 79 | **26** | **6** | **4** | **0** | **23** | 30 | NT | NT | 10 | 20 | 10 | 9 |
| P18 | 77 | **57** | **9** | **4** | **3** | 30 | 33 | 7 | 6 | 10 | 20 | 10 | 10 |
| P19 | NT | NT | NT | NT | NT | **6** | 32 | NT | NT | NT | **16** | 9 | NT |

| | | | | | | | | | | | | | |
|---|---|---|---|---|---|---|---|---|---|---|---|---|---|
| P20 | 78 | **14** | **3** | 6 | **2** | **24** | **19** | **4** | **2** | 10 | **17** | 10 | 7 |
| P21 | NT | **19** | **5** | **3** | **3** | 30 | **25** | **2** | 6 | 9 | 18 | 9 | 8 |
| P22 | NT | NT | NT | **0** | **0** | 16 | 35 | NT | NT | NT | NT | NT | NT |
| P23 | NT | NT | NT | **2** | NT | 27 | 30 | NT | NT | NT | NT | NT | NT |
| # Tested | 15 | 17 | 17 | 22 | 19 | 22 | 23 | 19 | 19 | 17 | 18 | 18 | 17 |
| # Impaired | 7 | 15 | 16 | 16 | 14 | 8 | 9 | 5 | 7 | 3 | 5 | 3 | 7 |

*Note.* Scores are number of correct responses unless otherwise specified. NT = unavailable for testing; TEA = Test of Everyday Attention, elevator counting subtest; VOSP = Visual Object and Space Processing battery. Category fluency corresponds to 8 categories (animals, fruit, birds, breeds of dog, household objects, tools, vehicles, types of boat). Letter fluency corresponds to F, A, S. Cut-offs for impairment correspond to two standard deviations below control mean performance, with impaired scores underlined and in bold. These are taken from control norms from respective tests manuals, unless otherwise specified (see below).

[a] Cut-offs taken from control testing at the University of York. Number of controls = 20.

*Supplementary Table 2. Patient performance on the Cambridge Semantic Battery and tests of semantic control.*

| | Semantic cognition composite score | Picture Naming | | Word-picture matching | CCT | | Ambiguity | | | | | | Synonym with distractors | | Object use | |
|---|---|---|---|---|---|---|---|---|---|---|---|---|---|---|---|---|
| | | No cues | With cues | | Word | Picture | Miscued dominant | Miscued subordinate | No cue dominant | No cue subordinate | Cued dominant | Cued subordinate | Strong distractor | Weak distractor | Alternative | Canonical |
| Max | - | 64 | 64 | 64 | 64 | 64 | 30 | 30 | 30 | 30 | 30 | 30 | 42 | 42 | 37 | 37 |
| Cut-off | - | 59 | - | 62.7 | 56.6 | 52.7 | 30 | 26.6 | 28.4 | 27.6 | 30 | 28.8 | 35.4 | 40.4 | 33.7 | 35.9 |
| Mean | - | 35.0 | 48.2 | 58.6 | 47.4 | 47.7 | 18.4 | 13.5 | 23.7 | 16.1 | 23.2 | 21.6 | 20.8 | 29.1 | 22.0 | 34.3 |
| P01 | .76 | **51** | NT | **50** | **54** | 54 | NT | NT | **26** | **23** | NT | NT | NT | NT | **24** | 35 |
| P02 | -.71 | **30** | NT | **54** | **41** | **46** | NT | NT | **19** | **10** | NT | NT | NT | NT | NT | NT |
| P03 | -.51 | **21** | NT | **46** | **42** | **44** | NT | NT | **21** | **13** | NT | NT | NT | NT | **12** | **30** |
| P04 | -2.62 | **5** | NT | **48** | **16** | **15** | **5** | **7** | **11** | **10** | **12** | **14** | **15** | **18** | **9** | **31** |
| P05 | -1.62 | **5** | NT | **50** | **33** | **13** | **19** | **9** | **23** | **10** | **24** | **17** | **18** | **34** | **13** | **31** |
| P06 | -.72 | **55** | NT | **60** | **39** | **36** | **18** | **10** | **23** | **13** | **22** | **22** | **19** | **24** | **24** | 37 |
| P07 | 1.30 | 62 | NT | 64 | 60 | 61 | NT | NT | 29 | **24** | NT | NT | **29** | **36** | **31** | 37 |
| P08 | .66 | **0** | 0 | **56** | **56** | 61 | NT | NT | **25** | **16** | NT | NT | **16** | **33** | **22** | 35 |
| P09 | .22 | **50** | 63 | 64 | **53** | 56 | **14** | **8** | **22** | **14** | **22** | **18** | **20** | **24** | **21** | 35 |
| P10 | -.72 | **19** | 58 | **60** | **29** | **45** | **13** | **14** | **24** | **14** | **19** | **20** | **13** | **29** | **14** | **29** |
| P11 | .73 | **3** | 10 | **52** | 57 | 54 | **21** | **18** | **27** | **19** | **23** | **24** | **30** | **31** | **22** | **33** |
| P12 | -.78 | 61 | 63 | **62** | **43** | **44** | **13** | **10** | **18** | **9** | **21** | **14** | **12** | **23** | **13** | **31** |
| P13 | .96 | **46** | 64 | 63 | **56** | 61 | **26** | 28 | **27** | **21** | **29** | **28** | 38 | **36** | **32** | 37 |
| P14 | .96 | **56** | 62 | 64 | 61 | 53 | **24** | **18** | **28** | **21** | **27** | **23** | **22** | **28** | **26** | 37 |
| P15 | -.98 | **1** | 3 | 63 | **39** | **31** | **12** | **7** | **22** | **11** | **23** | **25** | **15** | **25** | **14** | **32** |
| P16 | .24 | **50** | 63 | 63 | **48** | **51** | **27** | **16** | **25** | **18** | **27** | **25** | NT | NT | **27** | 37 |
| P17 | .52 | **50** | 64 | **62** | **52** | 57 | **19** | **15** | **26** | **17** | **23** | **28** | **23** | **30** | **22** | 35 |

| | | | | | | | | | | | | | | | | |
|---|---|---|---|---|---|---|---|---|---|---|---|---|---|---|---|---|
| P18 | 1.04 | 62 | 64 | **62** | 60 | 61 | **26** | **19** | **28** | **19** | **29** | **25** | **17** | **39** | **29** | 37 |
| P19 | .55 | NT | NT | 61 | **50** | 59 | NT | NT | **27** | **17** | NT | NT | NT | NT | **24** | **33** |
| P20 | .43 | 60 | 64 | **62** | 59 | **45** | **20** | **10** | **24** | **19** | **24** | **19** | **21** | **27** | 34 | 37 |
| P21 | .28 | **12** | NT | 64 | **48** | 54 | NT | NT | **23** | **19** | **NT** | **NT** | **25** | **29** | **27** | 37 |
| P22 | - | NT | NT | NT | **7** [a] | **10** [a] | NT | NT | NT | NT | NT | NT | NT | NT | NT | NT |
| P23 | - | NT | NT | NT | **13** [a] | **10** [a] | NT | NT | NT | NT | NT | NT | NT | NT | NT | NT |
| # Tested | - | 20 | 12 | 21 | 23 | 23 | 14 | 14 | 21 | 21 | 14 | 14 | 16 | 16 | 20 | 20 |
| # Impaired | - | 16 | - | 12 | 18 | 12 | 14 | 13 | 20 | 21 | 14 | 14 | 15 | 16 | 19 | 8 |

*Note*. Scores are number of correct. NT = unavailable for testing, CCT = Camel and Cactus Test. Cut-offs for impairment are taken from testing at the University of York and correspond to two standard deviations below mean control performance, with impaired scores underlined and in bold. Number of controls: CCT, Picture naming, and Word-picture matching = 10, Ambiguity task, Synonym with distractors, Object use = 8. Semantic composite score reflects regression scores derived from principal components analysis, including performance on CTT words, CCT pictures, and the Ambiguity task (no cue: dominant + subordinate). Lower composite scores reflect greater impairment.

[a] Patients P22 and P23 completed short versions of the CCT tasks, each comprising 25 trials. Cut-off for impairment for the word and picture versions of the task is 20.7 and 19.6, respectively. As these patients do not have scores for the long version of the CCT tasks or the Ambiguity task, they do not have semantic composite scores.

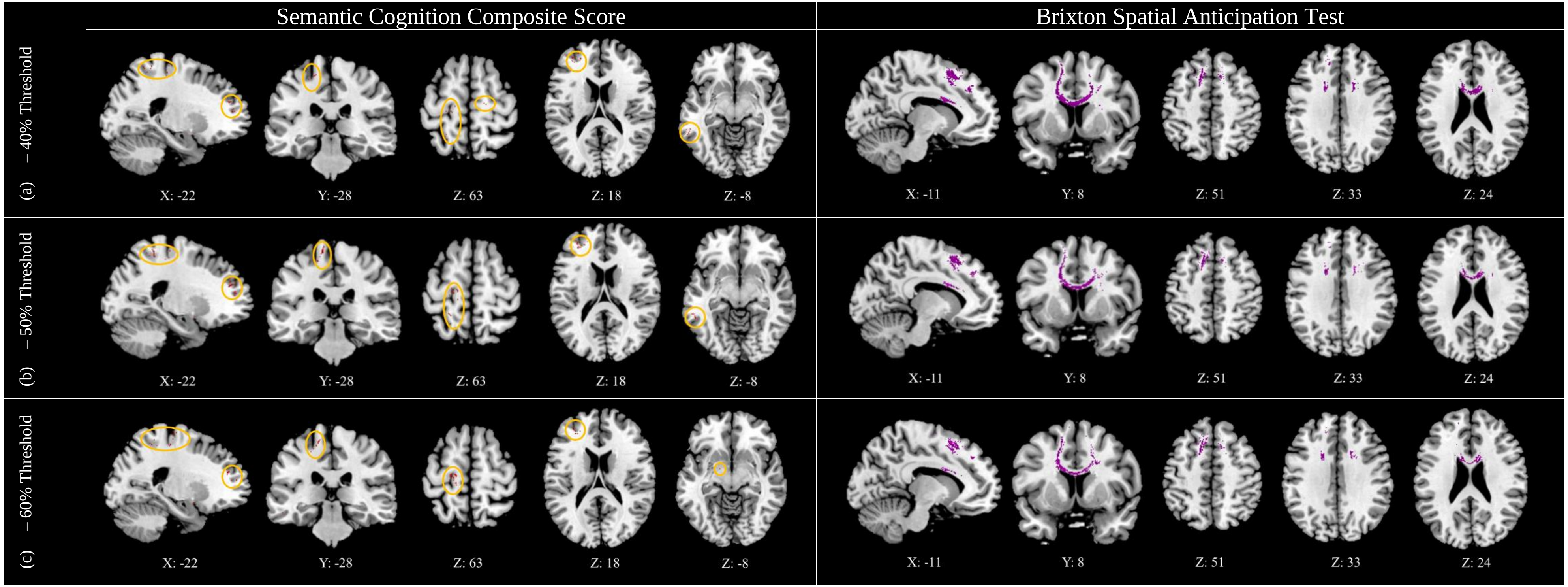


*Supplementary Figure 1. Structural disconnection symptom mapping at alternative thresholds for the likelihood of disconnection in the BCBtoolkit including at (a) 40%, (b) 50% (used in formal analysis), and (c) 60%, used to establish the stability of our structural disconnection analysis. Shown for clusters associated with lower semantic cognition composite scores (left) and lower scores on the Brixton Spatial Anticipation Test (right). Generated using non-parametric permutation tests in Randomise with threshold-free cluster enhancement. Highlighted voxels have a t-value of 2.6 or higher. Small clusters are highlighted in orange circles. 3D rendering generated in SurfIce. N = 20.*

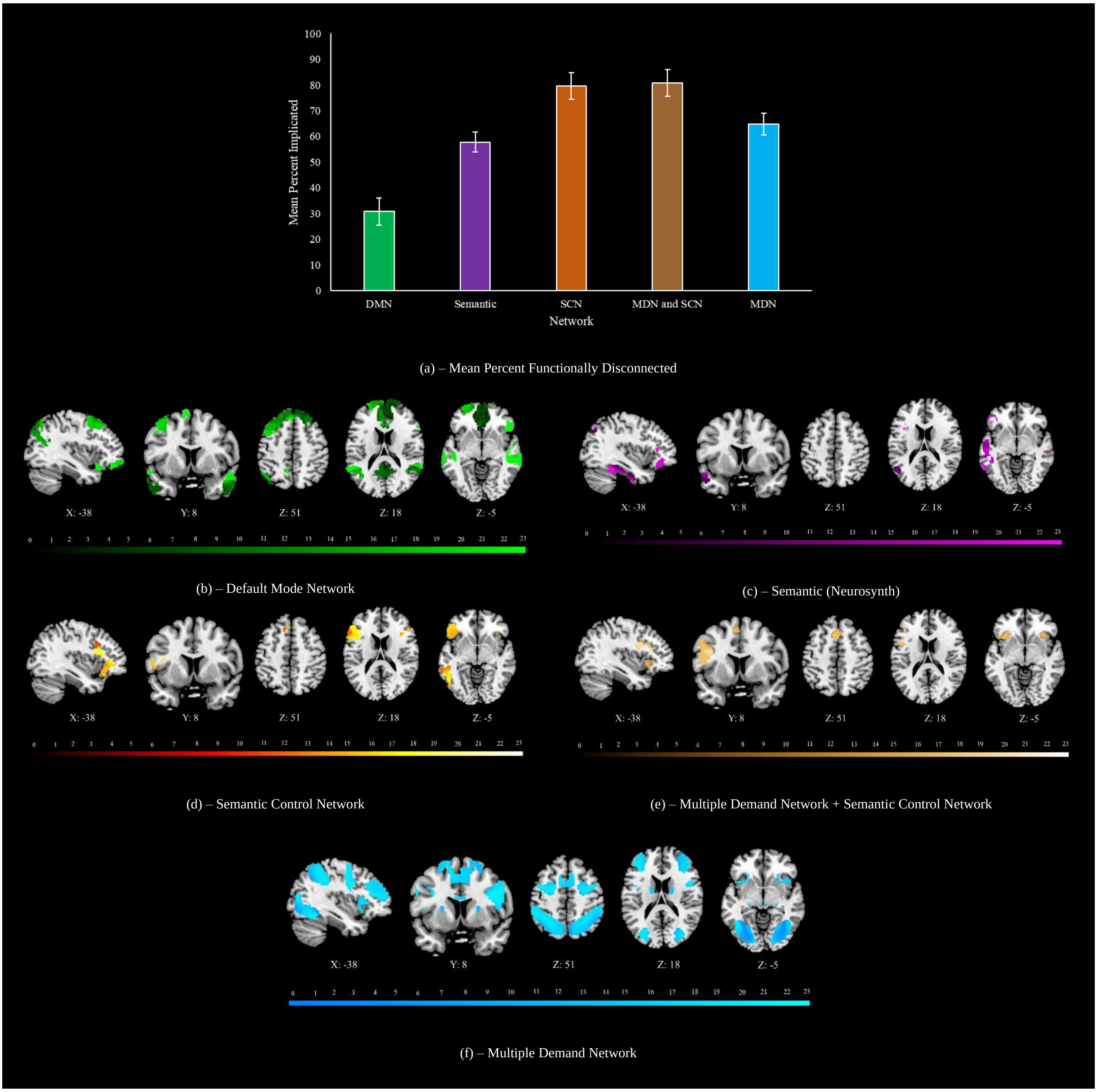


*Supplementary Figure 2. (a) The mean percent of each network of interest overlapping with patient functional disconnection maps, generated in CONN. DMN = default mode network, SCN = semantic control network, MDN = multiple demand network. Networks are visualised for (b) the DMN (Yeo et al., 2011), (c) a 'semantic' map (taken from a term-based meta-analysis from Neurosynth), (d) the SCN (Jackson, 2021), (e) areas common to the MDN and SCN, and (f) the MDN (Fedorenko et al., 2013). Keys under each map reflects the number of patients whose map overlap in a given voxel. N = 23.*

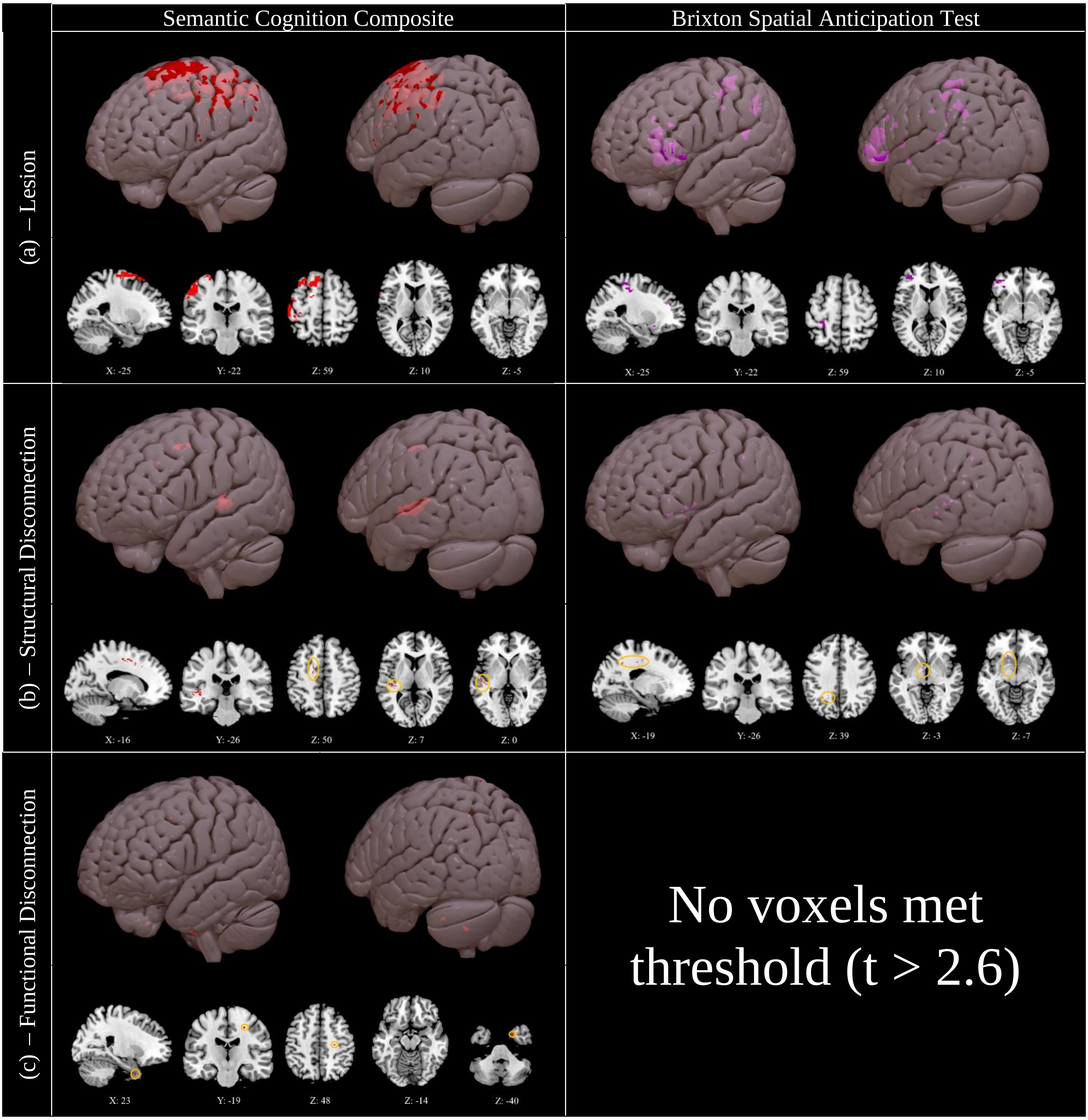


*Supplementary Figure 3. Voxels associated with higher semantic cognition composite scores (left) and better performance on the Brixton Spatial Anticipation Test (right), for (a) lesion, (b) structural disconnection, and (c) functional disconnection data. Generated using a non-parametric permutation tests in Randomise with threshold-free cluster enhancement. Highlighted voxels reflect those with a t-value of 2.6 or higher. Small clusters are highlighted in orange circles for visibility. 3D rendering generated in SurfIce. N = 20.*

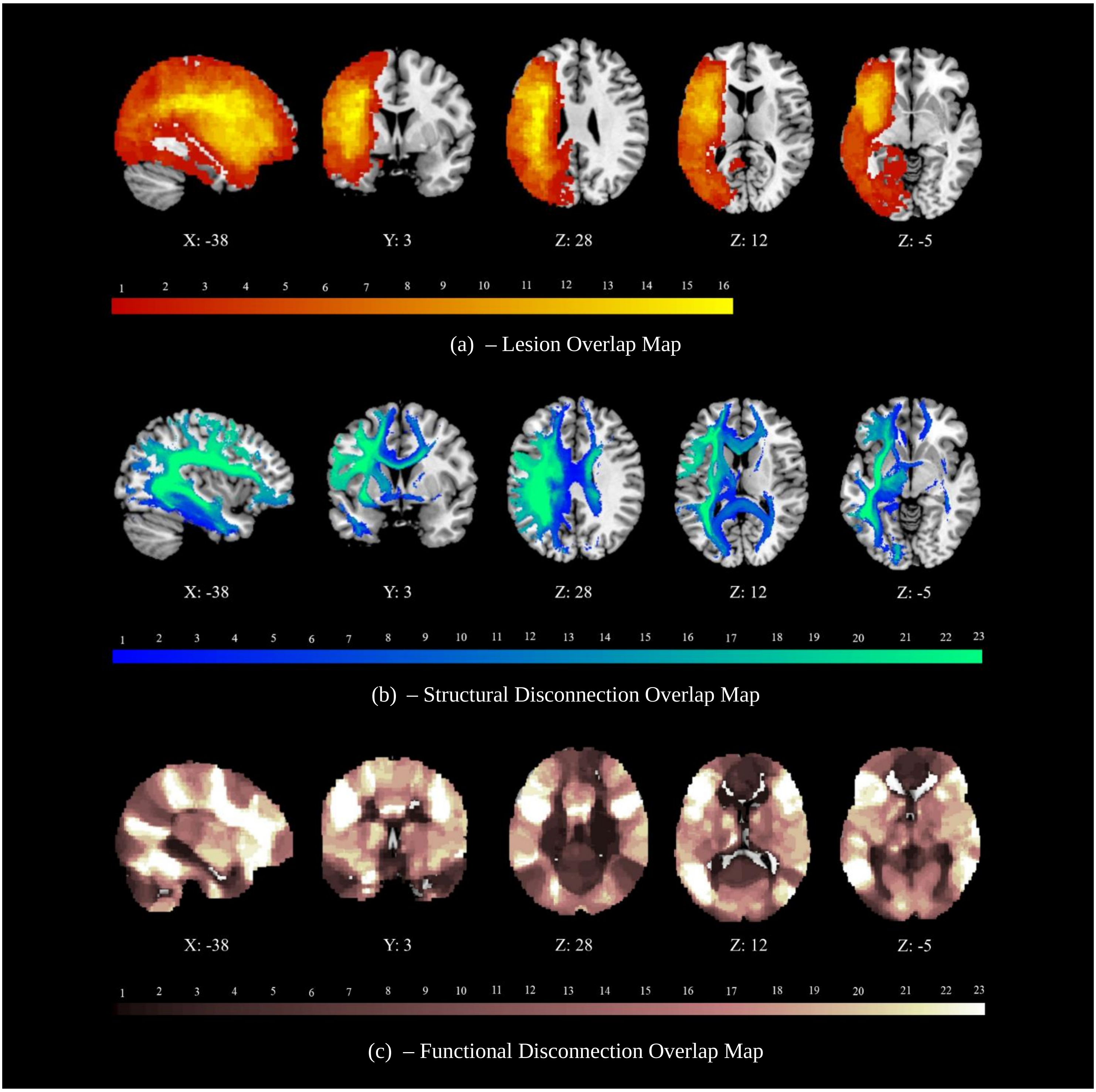


*Supplementary Figure 4. Unthresholded overlap maps for (a) lesion sites, (b) structural disconnection maps, generated using the BCB Toolkit, and (c) functional disconnection maps, generated using CONN. N = 23.*

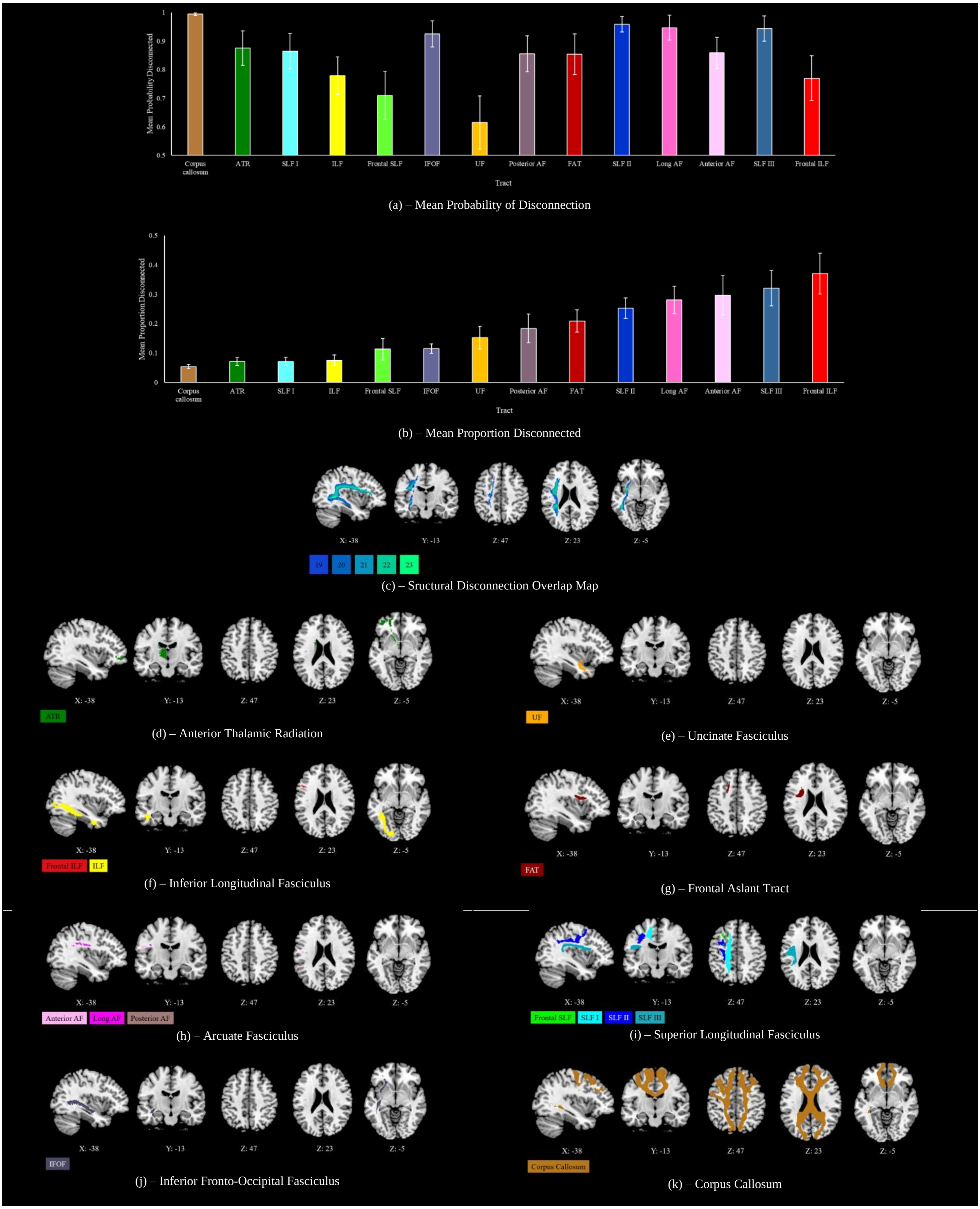
Mean Probability Disconnected
1
0.9
0.8
0.7
0.6
0.5
Corpus callosum
ATR
SLF I
ILF
Frontal SLF
IFOF
UF
Posterior AF
FAT
SLF II
Long AF
Anterior AF
SLF III
Frontal ILF
Tract
(a) – Mean Probability of Disconnection
Mean Proportion Disconnected
0.5
0.4
0.3
0.2
0.1
0
Corpus callosum
ATR
SLF I
ILF
Frontal SLF
IFOF
UF
Posterior AF
FAT
SLF II
Long AF
Anterior AF
SLF III
Frontal ILF
Tract
(b) – Mean Proportion Disconnected
X: -38
Y: -13
Z: 47
Z: 23
Z: -5
19
20
21
22
23
(c) – Sructural Disconnection Overlap Map
ATR
(d) – Anterior Thalamic Radiation
UF
(e) – Uncinate Fasciculus
Frontal ILF
ILF
(f) – Inferior Longitudinal Fasciculus
FAT
(g) – Frontal Aslant Tract
Anterior AF
Long AF
Posterior AF
(h) – Arcuate Fasciculus
Frontal SLF
SLF I
SLF II
SLF III
(i) – Superior Longitudinal Fasciculus
IFOF
(j) – Inferior Fronto-Occipital Fasciculus
Corpus Callosum
(k) – Corpus Callosum

*Supplementary Figure 5. (a) The mean probability of a given white matter tract being disconnected across the sample, and (b) the mean proportion disconnected. Generated using the Tractotron component of the BCB Toolkit (Foulon et al., 2018). Error bars reflect standard error of the mean. ATR = Anterior thalamic radiation, IFOF = Inferior fronto-occipital fasciculus, UF = Uncincate fasciculus, SLF = Superior longitudinal fasciculus, ILF = Inferior longitudinal fasciculus, AF = Arcuate fasciculus, FAT = Frontal aslant tract. (c) Structural disconnection overlap map, thresholded at 19 cases. White matter tracts are taken from Tractotron and thresholded at 0.95, including the (d) ATR, (e) UF, (f) ILF, (g) FAT, (h) AF, (i) SLF, (j) IFOF, and (k) corpus callosum. All tracts are confined to the left hemisphere. N = 23.*

*Supplementary Table 3. Main effects of network followed by Wilcoxon contrasts comparing the extent of both lesion and functional disconnection (percent of network impacted) between all functional networks of interest.*

| | | | | | |
|---|---|---|---|---|---|
| *Lesion* | | | | | |
| Network main effect | **F(2.2, 47.7) = 10.1, *p* < .001, $\eta_p^2$ = .32*** | | | | |
| | DMN | Semantic | SCN | SCN & MDN | MDN |
| DMN | | | | | |
| Semantic | Z = -2.4, *p* = .186 | | | | |
| SCN | **Z = -3.2, *p* = .015*** | Z = -2.5, *p* = .137 | | | |
| SCN & MDN | **Z = -3.5, *p* = .005*** | Z = -1.9, *p* = .569 | Z = -1.3, *p* > 1 | | |
| MDN | **Z = -4.2, *p* < .001*** | Z = -0.7, *p* > 1 | Z = -2.1, *p* = .333 | Z = -2.1, *p* = 358 | |
| *Functional Disconnection* | | | | | |
| Network main effect | **F(1.7, 38.2) = 27.2, *p* < .001, $\eta_p^2$ = .55*** | | | | |
| | DMN | Semantic | SCN | SCN & MDN | MDN |
| DMN | | | | | |
| Semantic | **Z = -3.9, *p* < .001*** | | | | |
| SCN | **Z = -3.9, *p* < .001*** | **Z = -3.3, *p* = .008*** | | | |
| SCN & MDN | **Z = -3.8, *p* = .002*** | **Z = -3.1, *p* = .019*** | Z = -1.0, *p* > 1 | | |
| MDN | **Z = -3.0, *p* = .030*** | Z = -1.8, *p* = .727 | Z = -2.5, *p* = .138 | **Z = -3.1, *p* = .019*** | |

Note: Non-parametric contrasts reported due to violation of the normality assumption. Results in both sections Bonferroni corrected for ten comparisons. * = significant result. N = 23.

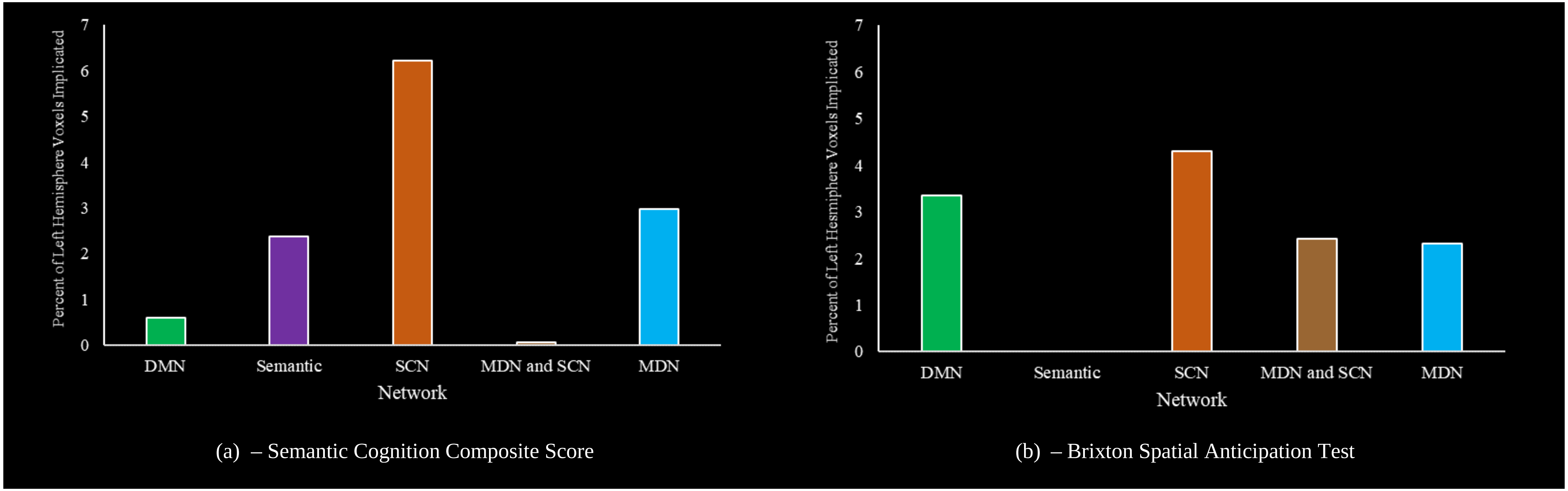


*Supplementary Figure 6. The percentage of voxels in each network of interest, restricted to the left hemisphere, implicated in the group level lesion-symptom mapping output for (a) the semantic cognition composite score and (b) the Brixton Spatial Anticipation Test. DMN = default mode network, SCN = semantic control network, MDN = multiple demand network.*[1]

[1] Note that these percentages will be impacted by differences in the relative size of each network. The number of voxels implicated over the total size of the respective number of voxels in each network for the Semantic Cognition Composite Score is: DMN: 82/13,618, Semantic: 84/3,549, SCN: 220/3,538, MDN & SCN: 1/1,777, MDN: 380/12,731. For the Brixton Spatial Anticipation Test, it's: DMN: 457/13,618, Semantic: 0/3,549, SCN: 152/3,538, MDN & SCN: 43/1,777, MDN: 296/12,731. Due to these differences in size, comparisons for a give network between graphs will be most informative.
